# Time cells emerge early in learning and encode stimulus modality past task requirements

**DOI:** 10.1101/2024.07.28.605458

**Authors:** Soumya Bhattacharjee, Hrishikesh Nambisan, Upinder S. Bhalla

## Abstract

Hippocampal neurons represent multiple dimensions of stimulus and behavioural context, including time. It is unclear how soon time-encoding cells emerge during learning, and if they additionally encode context in a manner specific to behavioural paradigm. We investigated simultaneous time and context encoding using 2-photon calcium imaging of mouse hippocampus during a trace eyeblink conditioning (TEC) task with sound and light stimuli. We find that the fraction of time cells is independent of learning and of stimulus modality. Only 13% of cells retain time-encoding on successive days, but persisters remain active within their original stimulus and post-stimulus epochs. Finally, we show that 60% of modality-specific time-encoding cells are active after the stimulus period, but modality-agnostic time cells are rare post-stimulus. Thus, compared to other paradigms, time cells in TEC have distinct learning and turnover properties, and exhibit sustained coding of stimulus modality and time which may subserve associations with subsequent events.

## Introduction

Temporal context differentiates episodic memory from semantic or procedural ones (Tulving, 1983, 1972; Tulving and Markowitsch, 1998) and is thought to be underpinned by cells which fire in stimulus free time-points between salient events (Pastalkova et al., 2008). These cells, termed as time cells have been reported in working memory paradigms such as delayed nonmatch to sample (DNMS) (MacDonald et al., 2011), and have been shown not to be sensitive to location (Kraus et al., 2013). While these studies observed time cells in the seconds to tens of seconds range, time cells also occur in sub-second time-scales (Modi et al., 2014). There are suggestive similarities between place and time cells in the hippocampus (Eichenbaum, 2014). Both place cells and time cells show sparse encoding (McNaughton and Morris, 1987; MacDonald et al., 2013), theta phase precession (Skaggs et al., 1996; Pastalkova et al., 2008), remapping (Muller and Kubie, 1987; MacDonald et al., 2011) and day-to-day populational drift (Ziv et al., 2013; Mau et al., 2018a)

Both place and time cells manifest as sequential activity during behaviour, however it is unclear if spontaneous cell sequences (Dragoi and Tonegawa, 2011; Villette et al., 2015) are signatures of place or time, or a more general manifestation of hippocampal predisposition for sequential activity relevant to a range of contexts

Context is important for place cells, and one of the goals of the current study was to explore if the same was the case for time cells. For example, directional place cells fire based on whether the animal is moving in a clockwise or counter clockwise direction in an open field or whether the animal is moving left or right in a 2D track (Gothard et al., 1996; Markus et al., 1995; McNaughton et al., 1983; Olton et al., 1978). Distinct ensembles of place cells have been found to be active during approach and return episodes near rewards or starting points (Weiner, Eichenbaum 1989; Gothard et al., 1996(a); Gothard et al., 1996(b)). These evidences along with observations that the hippocampus encodes various non-spatial variables (Aronov et al., 2017; Eichenbaum et al., 1987; Herzog et al., 2019; Park et al., 2020; Rita Morais Tavares et al., n.d.; Schuck and Niv, 2019; Taxidis et al., 2020; Wood et al., 2000) support the theory that the hippocampus doesn’t create a spatial map or temporal map but a cognitive map (Whittington et al., 2020, O’Keefe and Nadel, 1978; Tolman, 1948). In line with this theory, it would be expected that time cells also encode a broad spectrum of environmental features, with their dynamics varying across different temporal paradigms (Taxidis et al., 2020; Bigus et al., 2024). However, most current studies employ tasks such as DNMS and spatial alternation tasks (Pastalkova et al., 2008), single day TEC (Modi et al., 2014) and fear conditioning (Ahmed et al.,2020), all of which incorporate limited analysis of how additional context influences their dynamics.

Additionally, it is known that place cells emerge very rapidly in new environments (Hill, 1978) but very few studies look at time cell emergence during learning, a phenomenon which may differ based on task demands. Modi et al. 2014 reported single day time cell emergence, linked to learning. Similarly, Taxidis et al.2020 observed time cells from day one, with an increase in proportion over days, correlated with learning.

The question of learning trajectory of time cells motivated our design of a slower, multiday TEC learning paradigm. To explore the role of context in the form of stimulus modality for time cells over learning, we introduced multiple stimulus modalities in our TEC protocol. We imaged two-photon calcium activity of hippocampal area CA1 pyramidal cells as mice sequentially learnt a TEC protocol with two different modalities. Our findings reveal early emergence and continued presence of time cell sequences throughout the acquisition of learning, during both trace and post-stimulus periods. However, the dynamics of these cells remained agnostic to the animal’s learning state. Interestingly, we found time cells which encoded stimulus modality as well as ones which were agnostic to them. We suggest that time cells may encode additional context in many behavioural situations, and their continued activity past the behaviourally relevant window may provide a network mechanism for learning further associations.

## Results

Our overall design used 2-photon calcium imaging of CA1 pyramidal neuron activity during acquisition of TEC with two modalities. We trained on one of sound or light, swapped modalities when the animals learnt, and finally tested both within a session using interleaved trials. Below we describe the protocols, report basal network responses, and then follow time cells over learning. We examine how time cells behave over successive sessions, and finally analyze modality encoding by time cells during and after the stimulus period.

### Longitudinal Two photon imaging of dorsal CA1 during multiple TEC paradigms

We established a three stage TEC protocol (Figure 1 A i,ii,iii) in which animals first learnt to pair CS1-US (learning1) with an air-puff to the eye (Figure 1Ai, Supplementary Figure 1A). There were two possible CS1 stimuli: a 50ms long Blue LED flash or a 50 ms 3500Hz tone (Figure 1B). A total of 10 animals were used for awake, behaving *in vivo* recording. 6 out of 10 animals underwent all three stages of the training protocol. 3 animals were trained on the light stimulus and 3 on the sound stimulus for CS1 in learning1. These stimuli were swapped for CS2 in learning2. 4 out of 10 animals learnt the light-puff pairing but were not taken to further stages in the training sequence. In all cases, a 50ms puff of air to the eye was used as the US. There was a 250ms stimulus-free gap between Conditioned Stimulus (CS) offset and Unconditioned Stimulus (US) onset. Each day the animals underwent a single session consisting of 60 CS-US pairings (Figure 1 Ai). The criterion for a Conditioned Response(CR) was defined as an eye closure of at least 10% of a full eye closure, timed between the CS onset and US onset (Figure 1 C,D. Supplementary Figure 1Av). behaviour score was calculated for each day as % CR. Animals had to show 60% behaviour Score in a session to progress to the next phase. Probe trials (only CS, no US) were introduced randomly at a probability of 1 trial in 10.

**Figure 1:**
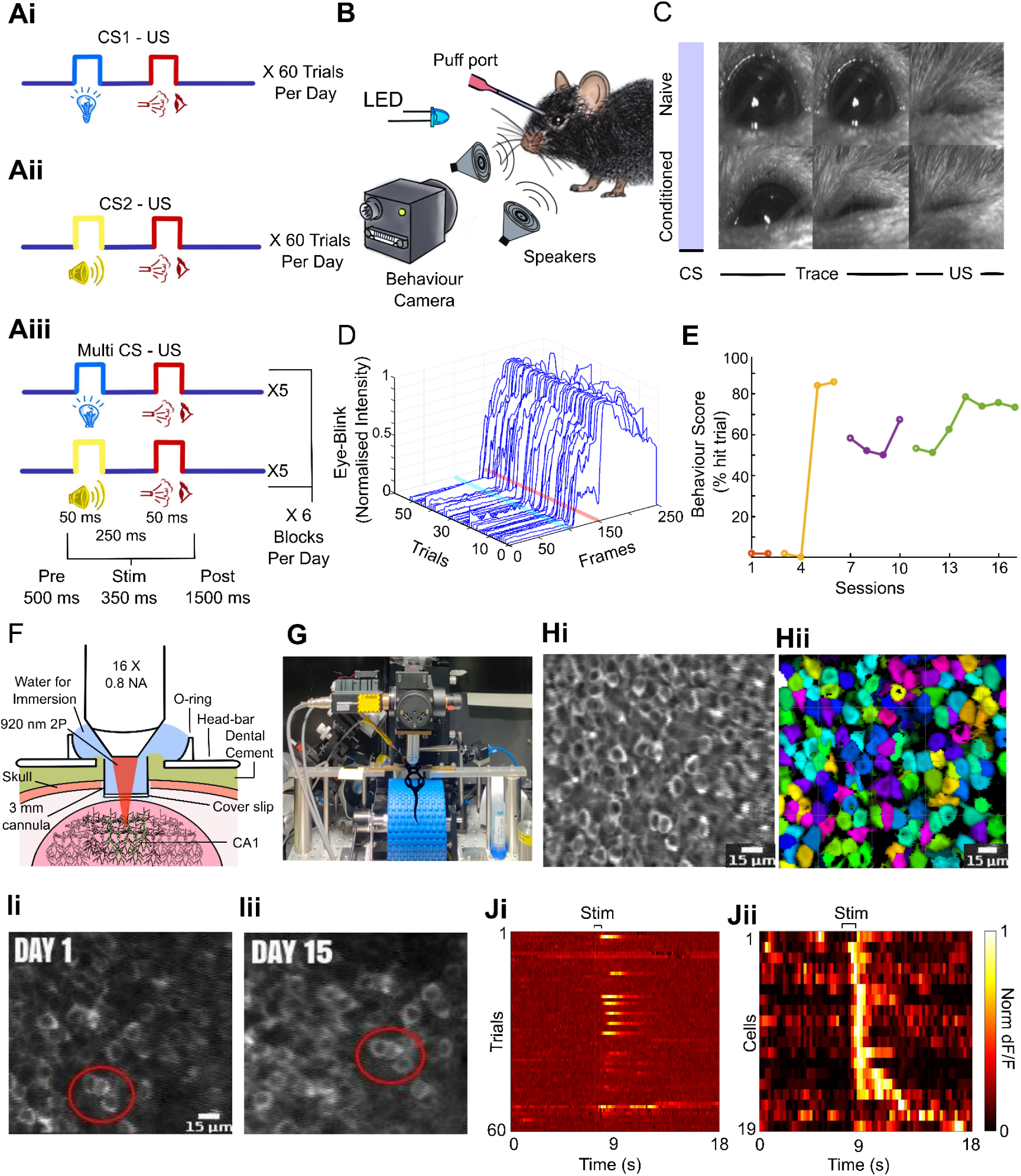
Neuronal and behavioural Dynamics in Trace Eyeblink Conditioning. **A i, ii, iii**: Trial structure of TEC protocol over the three phases of learning. There were 60 CS-US pairings per session. (i) Learning1, where animals learn the association between a conditioned stimulus (CS1, sound or light) and an unconditioned stimulus (US, air puff to eye). (ii) Learning2 where the CS is swapped (light for sound and vice versa). (iii) MultiCS phase where sound-US and light-US are presented in an interleaved manner. **B**: Schematic of the behaviour setup, depicting the two possible CS: a 50ms Blue LED flash and a 50 ms 3500Hz tone. US was delivered by a puff port and eye-blink responses were recorded by a 200 fps camera. **C**: Exemplar eye blink response of a naive(top) and trained (bottom) mouse. The blue bar denotes the time of CS. Trace and US time are marked at the bottom **D**: Waterfall plot shows all eye blinks in an exemplar session, timed between CS onset(blue bar) and US onset (red bar), across 60 trials. **E**: behavioural trajectory of mouse G142 over 17 days. Red: Rig habituation with no stimuli, yellow: Light-US pairing (learning1), purple: sound-US (learning2), green: multiCS phase. **F**: Schematic of chronic in vivo recording of Calcium activity, showing craniotomy, hippocampus, the positioning of a cylindrical cannula and a stainless steel headbar for head fixation under a microscope. **G**: Custom-built two-photon microscope setup used for imaging dorsal CA1 activity in head fixed mouse, running on a blue foam treadmill. **Hi** & **ii**: Automated ROI marking by Suite2p for extraction of fluorescent traces. **I i & ii**: Field of view imaged 15 days apart, illustrating the stability and repeatability of the imaging setup. **Ji**: Calcium activity of an exemplar time cell over 60 trials. Change in fluorescence signal over total signal (dF/F) was calculated using the 10th percentile of a cell’s fluorescent trace as a baseline, on per cell per trial basis. **Jii**: Raster plot showing trial averaged dF/F of 19 time cells in a session.

Learning1 of CS1-US took 5.1±1.1 days, mean±SEM. Next, CS modalities were swapped, i.e light to sound and vice versa (Supplementary Figure 1 Bii) in the CS2-US (learning 2) phase (Figure 1 Aii). Learning2 was typically faster (3.3±0.869, days mean±SEM. Supplementary Figure 1Bii). Once the animals reached and held the 60% criterion for a few more days (2±1.4, mean±SD), they were shifted to the third phase.

In the multiCS phase, sound-US and light-US were presented in the same session in an interleaved manner(Figure 1 Aiii). They were exposed to one block (5 trials) of Sound-Trace-US and then one block of Light-Trace-US. This continued for a total of 12 blocks: 6 for each condition, for the same total of 60 trials in each session. The last trial of each block was a probe trial where no US was delivered. Figure 1(E) shows the full behaviour journey of mouse G142 over 17 days, during which it went through all phases of the training protocol.

Simultaneously with the behaviour, we chronically recorded calcium activity from the pyramidal cells of the dorsal CA1 *in vivo*. Transgenic adult mice expressing Thy1-GCaMP6f+/- (GP5.17Dkim/J, Jackson Laboratories) underwent craniotomy and cortical aspiration (Dombeck et al., 2010) to expose the dorsal CA1. They were fitted with a 1.5mm long and 3mm diameter cylindrical cannula with a glass coverslip at the end (Figure 1F). Additionally a stainless steel headbar was implanted for head fixing under the microscope. Mice freely ran on a foam treadmill and were imaged using a custom built two photon microscope (Figure 1 F, G).

The recorded calcium data was motion corrected, classified into active regions of interest (ROIs) and fluorescent traces were extracted using the MATLAB version of Suite2p (Pachitariu et al. 2017) (Figure 1Hi & ii). Custom written MATLAB codes were used for analysis, but time cell classification used published Python/C++ codes (refs Ananth paper) called from MATLAB. dF/F was calculated using the 10th percentile of a cell’s fluorescent trace as a baseline,separately for each trial(Figure 1Ii).. For each session, the trial averaged dF/F was also calculated for establishing peri-stimulus time histogram (PSTH) heatmaps (Figure 1Jii).

Our recording spanned 163 sessions from 10 mice and yielded 11,543 cells (71±18 cells per session; mean±SD). Each mouse was recorded for 16±10 (mean±SD) days. Approximately the same field of view was imaged for all mice and sessions (Figure 1I i,ii). Multiday registration of the same cells were possible for 85 sessions from 5 mice.

### Network activity increases following stimulus delivery in a learning-independent manner

We next addressed the basic stimulus-driven responses of the hippocampal CA1 network. The hippocampus is known to receive multimodal sensory input (Acharya et al., 2016; Deshmukh and Bhalla, 2003; Ho et al., 2011; Komorowski et al., 2009; Liu and Otto, 2020). We divided each trial into 3 epochs: ∼1.7sec before the CS as pre-stimulus(pre), 350ms of CS, trace and US as stimulus(stim) and ∼1.7sec post US as post-stimulus(post). We defined calcium events as signals where the dF/F value was more than 2 standard deviations higher than the mean. We looked at trial averaged activity from 133 sessions over all stages of learning, from 10,383 cells.

Stimulus delivery elicited an increase in network activity, indicated by the frequency of calcium events (Figure 2A). We observed a significant rise in calcium event frequency from the pre-stimulus period to the stimulus period (Supplementary Figure 2C. p<0.0001, Wilcoxon matched-pairs signed rank test [WPT]), and continuing into the post-stimulus period (Supplementary Figure 2C. p<0.0001 WPT). Additionally, the post-stimulus increased activity occurred both in paired and probe trials, indicating that the increase was not just due to the presence of an aversive US (Supplementary Figure 2C. p<0.0001, WPT).

**Figure 2.**
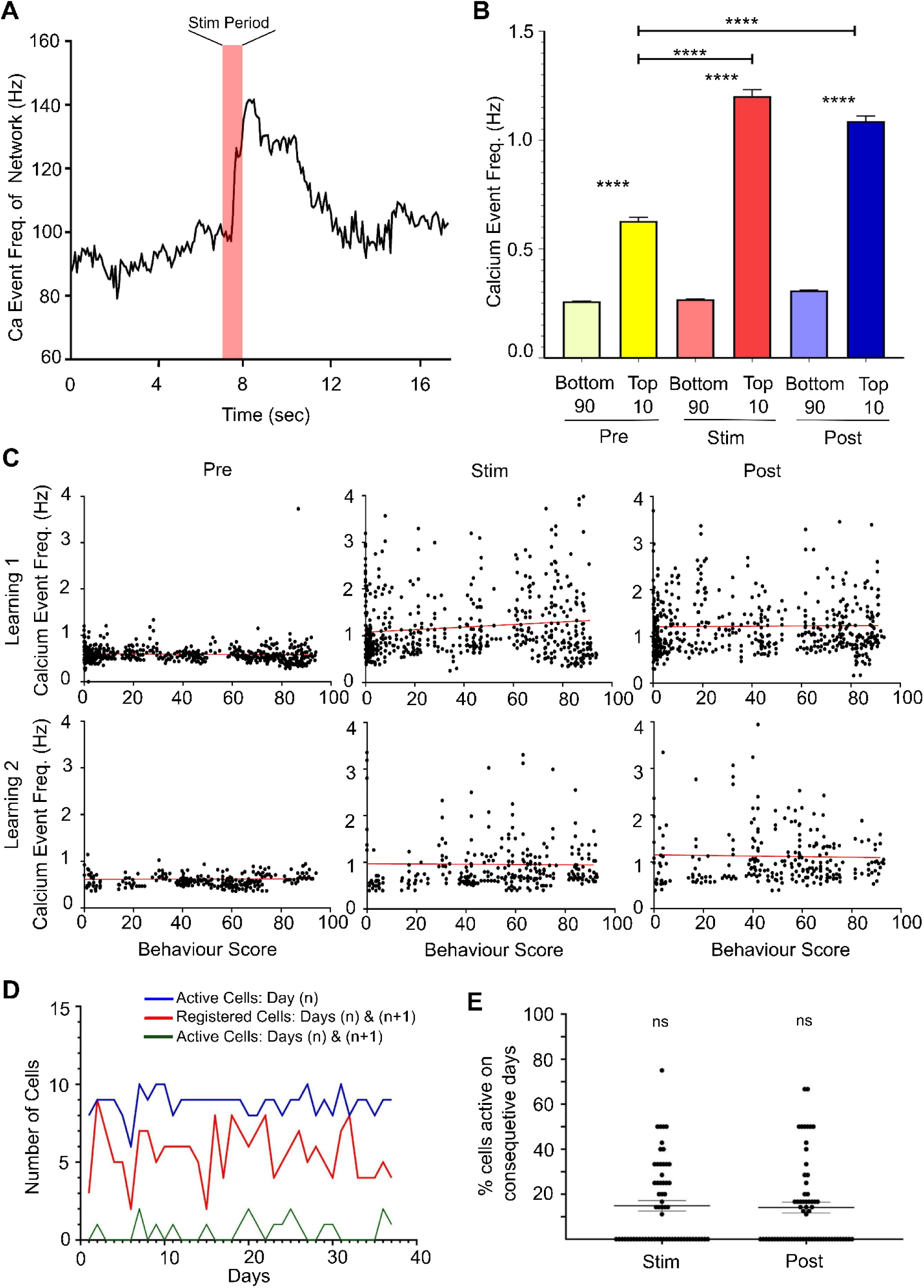
Stimulus-driven responses and learning effects on hippocampal CA1 network activity. **A:** Network activity increases upon stimulus delivery. Representative plot of calcium event frequency from all cells during a single session plotted as a function of time. The CS-US interval (“stim”) is indicated in pink. **B.** Calcium event frequencies are significantly higher in “active cells” (top 10% of most active cells) than the rest of the population, across all epochs. Pre: 0.66±0.02 Hz, Stim: 1.2±0.03 Hz, Post: 1.2±0.02 Hz vs. Other cells Pre: 0.27±0.001 Hz, Stim: 0.3±0.002 Hz, Post: 0.32±0.001 Hz. p<0.0001 for all comparisons between “Top 10” and “Bottom 90, Mann-Whitney test [MW]. Further, the top 10% most active cells showed significant increases in activity from the pre-stimulus phase (0.66±0.02 Hz) to the stimulus (1.2±0.03 Hz) and post-stimulus phases (1.2±0.02 Hz) (p<0.0001 for both comparisons, MW) **C.** Calcium activity of the top 10% active cells does not depend on learning. Per-cell, per-session activity is plotted against behavioural performance of nine animals for Learning1 and Learning2. No significant correlation was observed (linear regression F-test, p>0.05) except for small non-zero slope (0.0009) during Learning1 stim (F-test, p=0.003). **D.** Exemplar data from mouse G71 showing all active cells (blue curve), active cells that were registered (red curve) on the next day and cells which were active on both days (green curve). **E.** Activity status of a cell on a given day does not affect its chances of being active on the subsequent day. Comparison between proportion of cells active on consecutive days and population proportions (10%) showed no difference. Stim:13.56±2.157, Post:14.29±2.523.(One Sample Wilcoxon Text [OSWT], Stim p-value=0.4, Post p-value=0.52)

In every epoch we found that the total activity of the network was dominated by a small sub-population of cells (Figure 2B). This is expected since the hippocampus is known for sparse encoding (Jung and McNaughton, 1993; Skaggs et al., 1996). For each epoch, 10% cells had markedly higher activity rates (Mann-Whitney test [MW], p< 0.0001). This finding supports previous research, indicating that a small group of highly active cells, particularly in the Hippocampal CA1 region, are crucial in driving network responses (Agarwal et al., 2014; Senzai and Buzsáki, 2017; Treves and Rolls, 1994).

From one epoch to another, different sets of cells were active (Supplementary Figure 2F) and the level of activity increased from pre (0.628±0.017 Hz) to stim (1.20±0.030 Hz) and post (1.087±0.023 Hz) indicating the cells were about twice as active during and after stimulation compared to before (p<0.0001; MW, figure 2B).

We examined if learning affected neuronal activity in CA1. Prior studies suggest CA1 neuronal excitability increases during TEC (Berger et al., 1976; Christian and Thompson, 2003; McEchron et al., 2003, 1999; McEchron and Disterhoft, 1997; Moyer Jr. et al., 1996; Weiss et al., 1998). Figure 2C shows the activity of the top 10% active cells during learning1 and learning2 for each of the three epochs, plotted against behavioural performance of nine animals. Linear regression analysis found no significant correlation between behaviour scores and neuronal activity in either of the learning conditions (Figure 2C).

Using tracking of cells over multiple days (a total of 85 sessions over 5 animals) we found that an active cell is no more likely than any other cell to be active on the following day (p=0.38 and 0.52 for stim and post respectively. One Sample Wilcoxon test [OSWT]).(Figure 2D,E).

To summarize, we observed an increase in hippocampal CA1 network activity upon stimulus delivery. Heightened activity was maintained across both paired and probe trials. Notably, a small subset of highly active cells predominantly contributed to this increased activity. Activity did not significantly correlate with behavioural performance in learning phases, nor did a cell’s activity on one day predict its activity on subsequent days.

### Time cell fraction is independent of behavioural performance

Next we tested for time cell activity in hippocampal CA1 pyramidal neurons as a result of learning TEC. We assessed time-locking through precision (reduced peak firing time variability across trials) and hit rate (the proportion of trials with firing at a specified time), employing the Ridge-to-background (R2B) metric as defined by (Modi et al., 2014)(figure 3A). We compared each neuron’s peak trial-averaged calcium activity against that of a bootstrapped neuron—achieved by circular permutation of trial activities and averaging over 1000 cycles. An R2B score of 1 indicates timing accuracy equivalent to a randomized, bootstrapped neuron, and this was observed in 49.5% of neurons (figure 3B). Only neurons with an R2B score exceeding 2 were classified as time cells.

**Figure 3:**
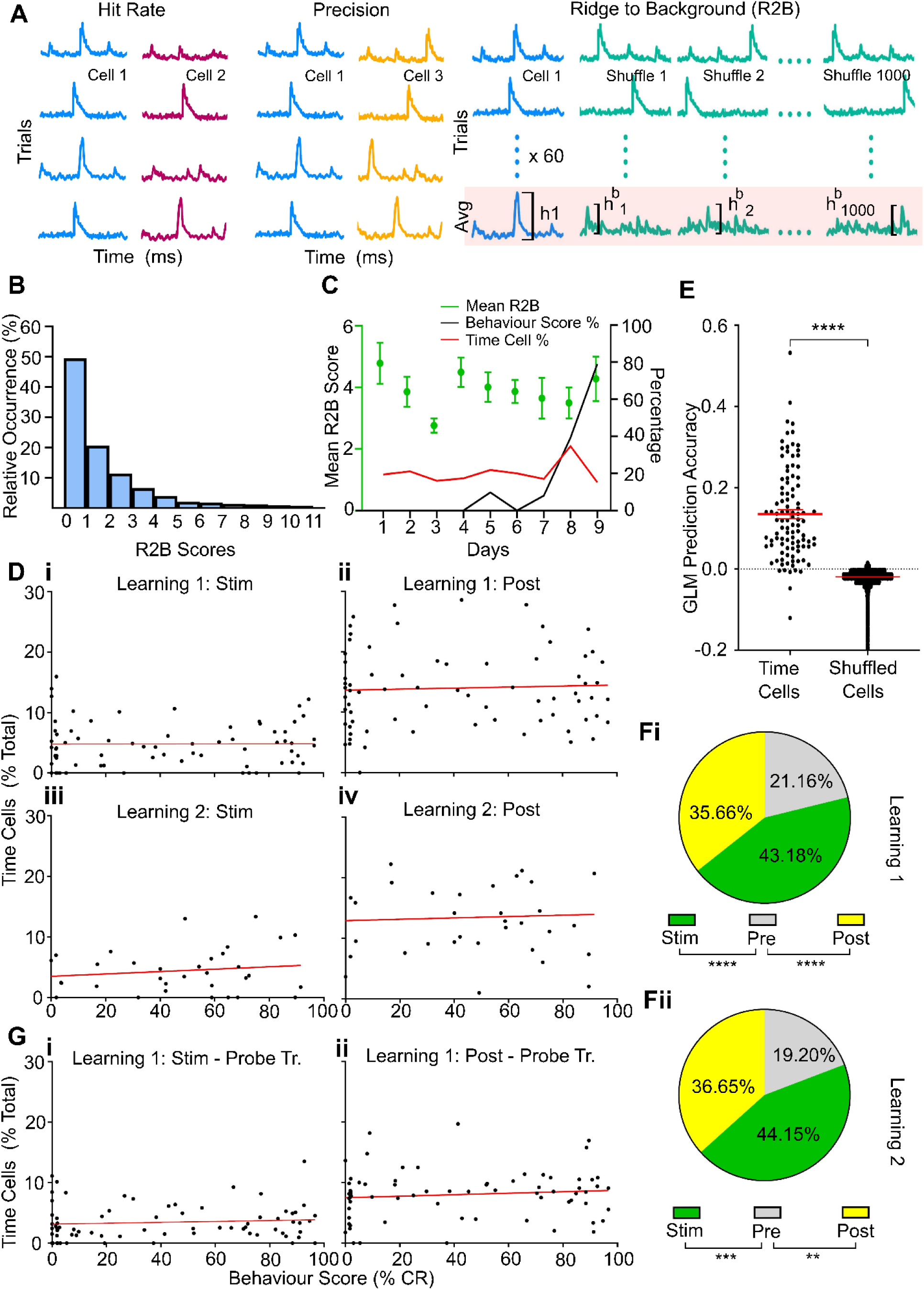
Learning Independent, time-locking in Hippocampal CA1 Neurons During TEC. A:Illustration of calculation of the Ridge-to-background (R2B) metric. It combines time cell precision and hit rate to obtain raw R2B for a neuron which is its peak, trial-averaged calcium activity. Bootstrapping is done by circularly permuting trials and then averaging. The R2B score is the ratio of the raw R2B to the mean of 1000 bootstrapped R2Bs. Cells with an R2B score above 2 are selected as time cells. B: Distribution of R2B scores across neurons. Around 70% did not qualify as time cells (R2B<2). C: Mean R2B scores of time cells for an exemplar animal (dots with whisker error bars) and proportion of time cells in population (red line) do not correlate with behavioural performance (solid black line) over successive days. D: Percentage of time cells do not change over learning stages. Each dot is the percentage of cells identified as time cells in a given session, data from all animals. Note that stimulus time=350ms vs post-stimulus period= 1.7s and percentage of time cells is over the entire window. Di: Mean percentage during the stimulus = 4.79±0.426% for CS1-US Dii: Mean percentage during post-stimulus=13.98±0.801% for CS1-US. Diii: Similarly, for CS2-US, during stimulus percentage=4.43±0.64%. Div: CS2-US post-stimulus percentage= 13.45±1.325%. Linear regression analysis showed no significant changes across learning stages (stimulus period slope: -0.0006±0.011; post-stimulus period slope: 0.0008±0.022 for CS1-US, and stimulus period slope: 0.019±0.64; post-stimulus period slope: -0.011±0.049 for CS2-US; all not-significant, F-test), indicating consistent time cell behaviour. E. A Generalized Linear Model trained on time cell data predicts time with significantly greater accuracy than a model trained with shuffled data (time cell:0.13±0.01, shuffled cells:-0.019±0.00004, p<0.0001;mean±SEM, MW) Fi, ii: Proportions of hippocampal time cells with peak activity for all the three epochs, showing a high degree of time cell activity in the post-stimulus period for both Learning1 (35.66%) and Learning2 (36.65%) phases, adjusted for the duration of each period. Both stim and post show significantly more time cells than the pre period (p<0.01 for all comparisons;Wilcoxon matched-pairs signed rank test [WPT]). Gi, Gii: Time cells are active in probe trials without US, in both stimulus and post-stimulus periods. Post-stimulus time cells in probe trials suggest a persistence of time encoding in the network beyond the stimulus period

Initially, in learning1, an average of 4.79±0.43% (mean±SEM) of neurons were classified as time cells active within the stimulus period, whereas 13.98±0.80% (mean±SEM) were time cells that belonged to the post-stimulus period. Surprisingly, the distribution of these time cells did not exhibit significant changes across different learning stages, including naive, active learning, and fully learned states. Figure 3C demonstrates this with an exemplar plot of mouse G141, showing Learning1 behaviour Score (black curve), time cell proportions (red curve) and mean time cell scores of the time cells (green dot and whisker) over days. Linear regression analysis for stimulus and post-stimulus period from all animals for Learning1 show no significant correlation between behaviour score and time cell proportions (F-test, p>0.05) (Figure 3Di and ii). This pattern persisted during Learning2, with 4.43±0.64% (mean±SEM) of cells recognized as time cells during the stimulus period (figure 3Diii) and 13.45±1.325% (mean±SEM) in the post-stimulus period (figure 3Div) and no significant correlation between behaviour score and time cell proportions (F-test, p>0.05). Together these findings show stability in proportions through the animal’s progression through learning stages.

We verified our R2B time cell detection method by training a generalized linear model (GLM) using data from identified time cells, to predict time (Figure 3E). As a control we trained a model on shuffled data from the same time cells. The GLM trained on time cell data performed significantly better than the shuffle control (time cell:0.13±0.01, shuffled cells:-0.019±0.00004, p<0.0001;mean±SEM, MW). We further validated the presence of time cells using a temporal information metric (Mau et al., 2018b) (Supplementary Figure 3).

In summary, we found consistent time cell proportions across learning stages. As considered in the discussion, this contrasts with previous findings on time cell emergence over the course of learning in tasks with stimulus-free intervals.

### Many time cells are tuned to the post-stimulus period

We found that a large percentage of hippocampal time cells peaked in activity during the post-stimulus period, which we monitored for 1.7 seconds (figure 3Fi and ii, 35.66% in the Learning1 and 36.65% for Learning2, normalized for epoch duration). The proportions of time cells in stimulus and post-stimulus periods were always significantly higher than in pre period (p<0.01 for all comparisons; WPT). This held true both for Learning1 and 2. To determine whether this prolonged activity was triggered by the US, we analyzed probe trials with only the CS and no US. Time cells were active in both the stimulus and post-stimulus periods of probe trials (Stim: 3.43±0.33, Post: 7.98±0.45; mean±SEM), suggesting a role in encoding time well past the behaviourally relevant duration of the TEC. Moreover, the presence of time cells was not correlated with behavioural scores, indicating that post-stimulus time cell activity is independent of task performance (Figure 3Gi and II. Slopes of 0.007±0.009 for stim and 0.012±0.012 for post, both not significant, F-test).

Thus, we found cells with time-encoding even after the stimulus period, and this extended activity was independent of stimulus modality and task performance.

### Most time cells turn over, but the persisters fire in consistent epochs

We investigated time cell reliability across multiple days by tracking cells over 85 sessions involving 5 animals. We focused on cells that maintained a reliability score of 2 or higher over both days (figure 4A).

**Figure 4:**
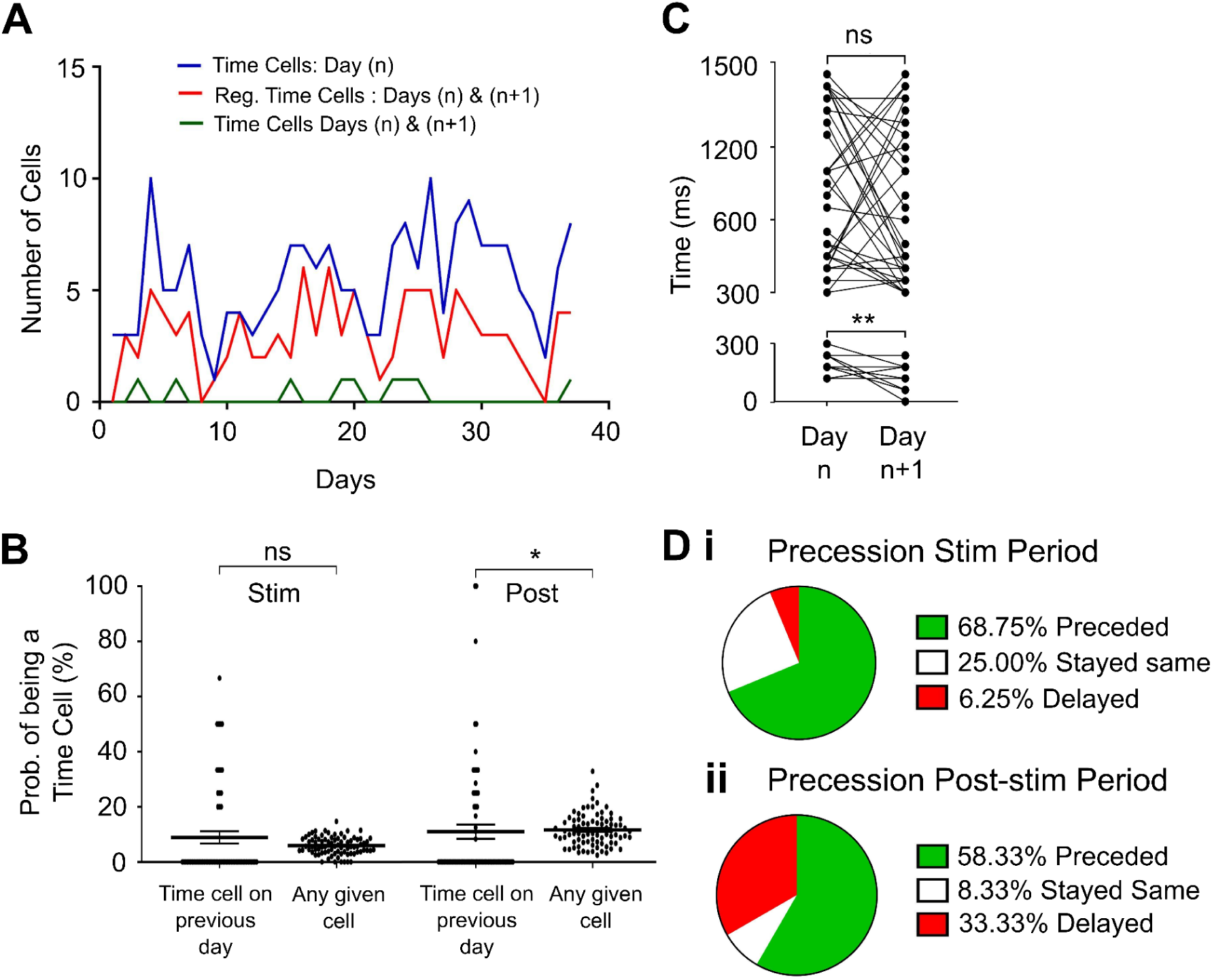
Day-to-Day Variability in Time Cell Identity with Stable Temporal Fields in Hippocampal CA1. A: Proportion of hippocampal CA1 cells identified as time cells on successive days in mouse G71. Blue: Classified Time Cells for Day(n). Red: Classified Time Cells for Day(n) which could be registered in the imaging on Day(n+1). Green: Time cells from Day(n), which were registered on Day(n+1) and retained time cell tuning. B: Percentage of time cells retaining their status from one day to the next (8.917±2.204% during stimulus and 10.98±2.575% post-stimulus). This percentage is comparable to the general probability of any given cell becoming a time cell (p=0.32 for stimulus period. p=0.03 for post-stimulus period, borderline significant, WPT). This indicates a high degree of remapping in the time cell population. C: Multi-day time cells firing in the stimulus period (<350 ms) always persist in the stimulus period across consecutive days (16 out of 16), and likewise those tuned to the post-stimulus period (>350 ms) remain there on the next day (36 out of 36). Further, time cells in the stimulus period showed a significant precession of time field (Day n: 216±12 ms vs. Day n+1: 111±18ms; p=0.0015, WPT) on the second day. Though the majority of cells in post-stimulus also showed precession, the population data was not statistically significant (p-value = 0.14, WPT). Di: Precession in time tuning during stimulus period. 68.75% of cells fired earlier on the following day. Dii: Precession in time tuning during post-stimulus period, 58.33% cells had earlier peak times on the following day

A cell that was time-tuned on a given day was unlikely to be time-tuned on the next day (∼91% dropouts for stimulus period and ∼89% for post-stimulus period) (figure 4B). The likelihood of a cell to remain time-tuned on the next day was similar to that of cells from the broader population becoming time tuned for the first time (8.9±2.2% during stimulus and 10.98±2.58% post-stimulus, vs 5.8±0.4% mean likelihood of any cell from the population being time-tuned during stimulus period and 11.6±0.7% during post-stimulus period). This difference did not show statistical significance during the stimulus period (p-value=0.32; WPT) and a borderline significance in the post-stimulus period (p-value=0.03; WPT). We do not ascribe any biological significance to it owing to the small difference in probabilities (10.98% vs 11.6%).

Despite the day-to-day turnover of most time cells, those that persisted across consecutive days (n=52) showed very consistent firing epochs, with stimulus period cells always firing during the stimulus period on the next day, and similarly for post-stimulus period cells (figure 4C). Interestingly, there was a statistically significant precession in firing time on the following day during the stimulus period for 68.75% of cells (Day n: 216±12 ms vs. Day n+1: 111±18ms; p<0.01, WPT, figure 4Di). While a shift towards earlier firing times was also noted in 58.3% of post-stimulus period cells, the change did not reach statistical significance(p-value=0.14, WPT).

Our results indicate a dynamic remapping of the time cell population day-to-day, rather than a stable population of time cells. The degree of remapping we observe is substantially higher than what has been reported in previous time cell studies (Mau et al., 2018b; Taxidis et al., 2020), as we consider in the discussion. Despite this day-to-day remapping, those cells that do persist show consistent firing epochs. Many time cells display a significant precession in peak firing times the following day, especially during the stimulus period.

### Transient network reliability marks learning onset

We next investigated whether there was a network correlate, as opposed to individual time cell correlates of learning. To do this, we introduced a “network reliability” metric, defined as the sum of reliability scores from cells scoring above 2, weighted by the time cell proportion in the network. This measure was predicated on the notion that a network with high temporal reliability would either have an increase in time cell proportion or have time cells with better time tuning, i.e. higher reliability score, or some combination of both.

This metric showed a correlation with learning in the stimulus period of Learning1 (Figure 5A). We found that network reliability remained low across 58 days from 9 mice, except for a noticeable and statistically significant spike on one day (figure 5B p<0.0001, OSWT). Remarkably, for 8 out of 9 mice that learnt the CS1-US protocol, network reliability peaked either just before or on the day behavioural performance met the 60% conditioned response criterion. Moreover, behaviour scores rose substantially after this surge in network reliability (figure 5C p<0.05, MW). On average the network reliability peaked 3±1.2 days before animals hit behaviour criteria (figure 5D. p<0.05 Paired t test, figure 5E). As we consider in the discussion, the pattern of network peak preceding behaviour peak was seen only in Learning1 in the stimulus period.

**Figure 5:**
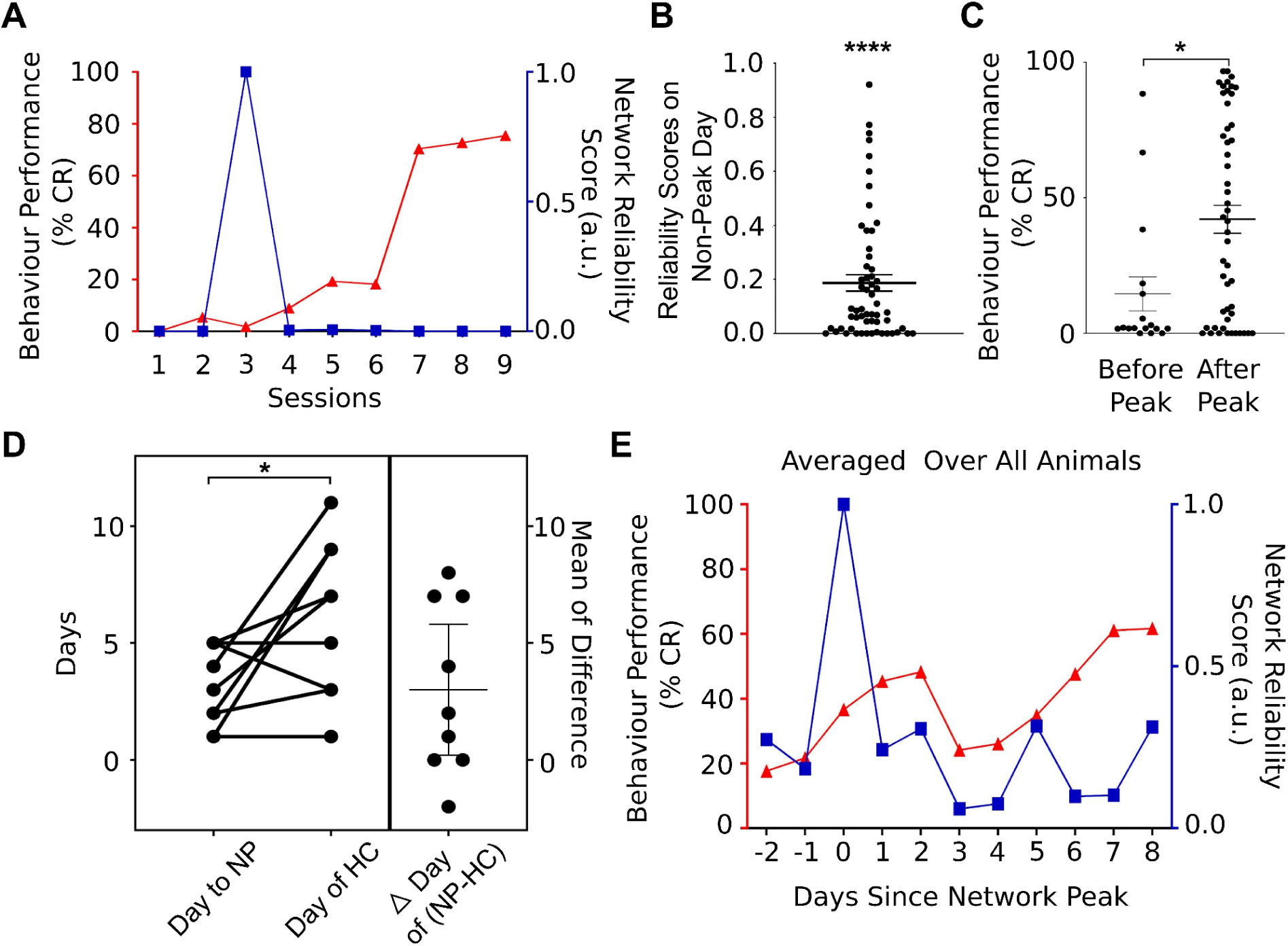
Transient Network Reliability Peaks as a Marker of Learning Onset. A: Network reliability metric illustrated for mouse G394 related to behavioural performance. Network reliability remains low except for a spike on Session 3, which precedes the improvement in behavioural performance.. B: Network reliability scores on non-peak days (0.186±0.03) significantly differ from those on peak days, normalized to 1 (p<0.0001, OSWT), suggesting a shift in network dynamics correlated with learning progress. C: behavioural scores rise following the peak in network reliability. behaviour Score (%Conditioned Response) before (14.55±6.282) and after (42.05±5.196) the peak in network reliability (p<0.05, MW) D: Network reliability peak (NP) occurs on or before the behaviour hitting criterion (HC) of 60%. Left: Paired data for each animal day of NP and day of HC. Right: Delay from NP to HC. NP typically precedes HC by 3±1.2 days (p<0.05, Paired t-test). 5E: Averaged peak network activity data across all mice, with peak days aligned to zero, showing that behaviour score rises following NP.

### Distinct Temporal Dynamics of Modality-Specific and Modality-Agnostic Time Cells in multiCS Sessions

Finally, we asked if time cell information was independent or correlated with modality information. To do so we utilized the multiCS phase of behaviour, during which the animals had to recall both stimulus modalities in an interleaved manner. This allowed us to contrast cell responses within an individual session. We utilized only those sessions in which behaviour performances exceeded 50%. In all, 6 scores were omitted out of a total of 58; 3 of the omitted scores were from the same animal.

We found some cells kept time irrespective of modality while others and some were unique to each condition (figure 6A. Modality-agnostic: 5.08±0.77%, total sound-US: 15.75±1.21%, total light-US: 16.64±1.313%). If these numbers were obtained from a uniform distribution, we would expect:

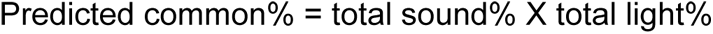

**Figure 6:**
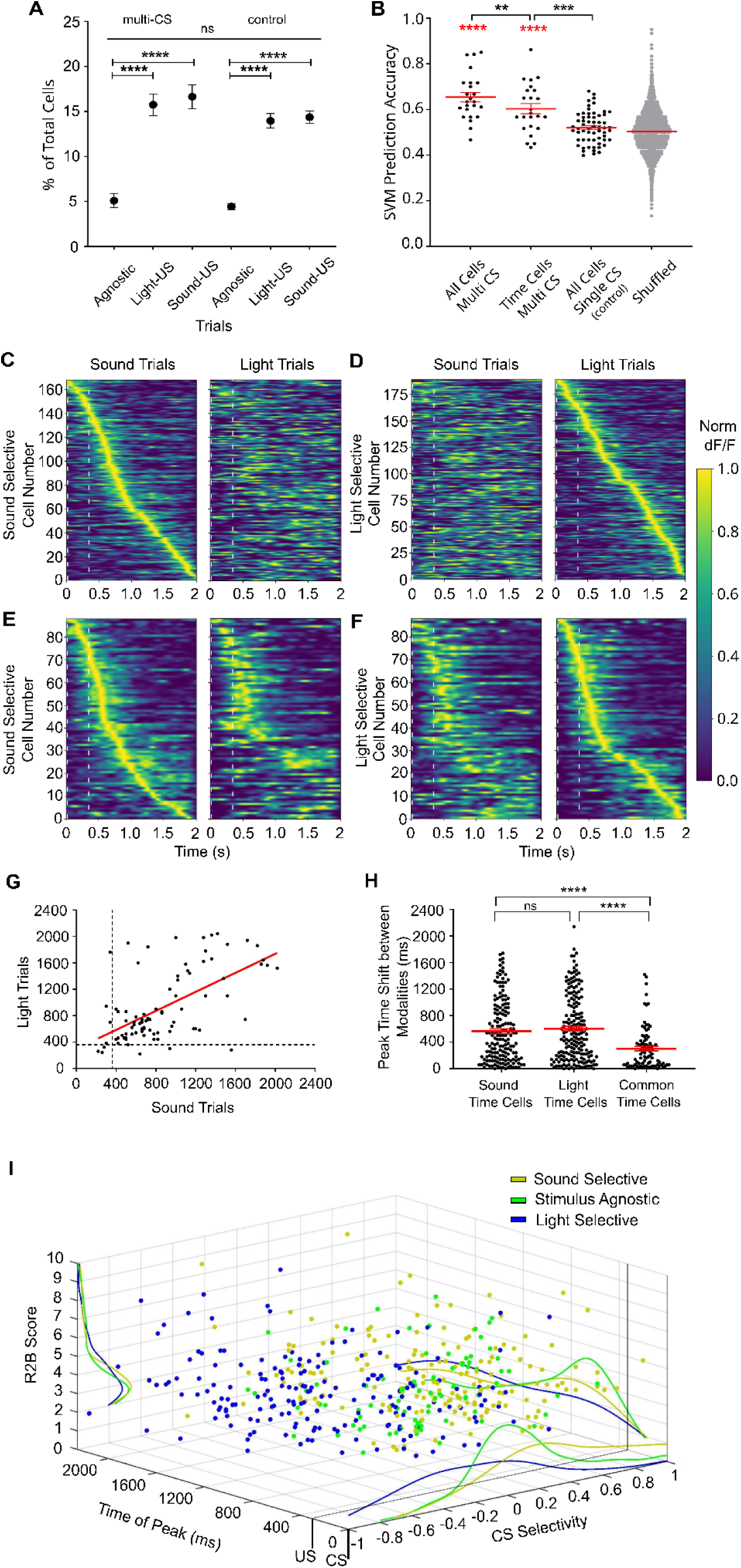
Distinct Temporal Dynamics of Modality-Specific and Modality-agnostic Time Cells in multiCS Sessions. **A**: Proportions of time cells which are either responsive in sound-US trial (15.75±1.2%), light-US trial (16.64±1.3%) or are modality-agnostic, i.e. responds in both conditions (5.08±0.77%). modality-agnostic time cells are significantly lesser in proportion than the other two populations (p<0.0001, MW) modality-agnostic time cells occur more frequently than those predicted by the product of likelihood of single-modality time cells (2.94%±0.38). For control comparison, single-CS-trials were subdivided into interleaved blocks to simulate the modality division of multiCS sessions (set1: 13.95±0.80%, set2: 14.37±0.70%, common: 4.43±0.34%). There was no significant difference observed between multiCS and Control (p=0.98,chi-square test). **B**: SVM prediction of modality from the activity of different categories of time cells. SVMs were trained on all cells from multiCS sessions,only time cells from multiCS sessions, all cells from singleCS sessions as control and modality shuffled (bootstrapped) multiCS data. SVMs trained on multiCS data - both all cells (0.65±0.02) and time cells (0.6±0.02) performed better than singleCS control (0.51±0.008) and bootstrapped data (0.5±0.005) which performed at chance level (p<0.0001 for all comparisons (red stars), MW). All cells have slightly more information about stimulus modality than just time cells (p=0.001, WPT). **C-F**: Heatmaps of normalized trial-averaged firing of time cells, pooled across all mice and multiCS sessions. As different sessions used slightly different frame-rates, each row was rescaled to time units by convolution with a 300-ms width Gaussian. Time cells were classified into three subtypes: sound-specific, light-specific, and modality-agnostic time cells (reliable to both CSs). C: Sound-specific time cells in multiCS sessions, sorted by time of peak in sound trials, during (Ci) sound trials, and (Cii) light trials. Sound-sorted sound trials form a clear sequence while the sound-sorted light trials do not. B: Light-specific time cells sorted by time of peak in light trials, during (Di) sound trials, and (Dii) light trials. E: modality-agnostic time cells sorted by time of peak in sound trials, during (Ei) sound trials and (Eii) light trials. F: modality-agnostic time cells sorted by time of peak in light trials, during (Fi) sound trials and (Fii) light trials. In the modality-agnostic cells there is significant residual time-ordering of cells even when sorted by the other modality. Also note that only 44% of modality-agnostic time cells are active after 0.8 seconds (as compared to 70% and 62% for light and sound-specific time cells) **G:** Stimulus agonistic time cells maintain time of peak firing between sound and light trials. (Linear regression R squared value = 0.36. Slope significantly different from zero; p<0.0001, F test) **H:** Absolute difference in peak times between sound and light trials are higher for sound and light-specific time cells as compared to modality-agnostic time cells (sound-specific:578±36ms, light-specific:604± 36ms, modality-agnostic: 300±34ms; mean±SEM. p<0.0001 for both comparisons vs modality-agnostic cells and p=0.51 for sound-specific vs light-specific; MW). **I:** Represents a cell’s location in 3D space, defined by its CS selectivity (x-axis, -1 for purely light to 1 for purely sound), peak firing time in a trial (y-axis), and time cell score (z-axis). The three categories of time cells (sound-specific, light-specific and modality-agnostic) as determined by their CS selectivity do not show any difference in level of time tuning (p=0.59, KWT) but show variation in time of peak activity. modality-agnostic “pure” time cells prefer to fire in and around the stimulus period whereas the modality-specific context encoding time cells fire throughout the stimulus as well as post-specific period.

Applying this calculation we obtained predicted common% = 2.62±0.29%. The predicted common fraction is significantly smaller than the observed common fraction (p=0.0014, Unpaired t test).

We conducted a control analysis using data from a single-modality CS-US session but subdivided into dummy interleaved CS1 and CS2 trials as if it were a multiCS session. The control distribution mirrored the multiCS time cell distribution: there were cells common to both sets, as well as unique cells within each set (figure 6A common: 4.43±0.34%, set1: 13.95±0.8%, set2: 14.37±0.7%). There was no significant difference between these two distributions (p=0.98, chi sq test).

Can one classify modality from time cell activity using an SVM (Fig 6B)? We used four datasets for training: all cells in multiCS sessions; only time cells from multiCS sessions; all cells from dummy interleaved Single CS sessions as control; modality shuffled MultiCS data (SVM accuracy for multiCS all cells=0.65±0.02, multiCS time cells=0.6±0.02, singleCS=0.51±0.008, shuffled=0.5±0.0005. Mean±SEM). SVM trained on time cells from multiCS sessions performed significantly better than the SingleCS control (p<0.0001, MW) and from shuffled multiCS data (p<0.0001, MW). Thus in multiCS sessions the CA1 time cells carry information about the CS modality. Interestingly, SVM trained on all cells’ data performed better than just time cells’ (p=0.001,WPT) implying modality-specific data in time cells is no more than the general population of CA1 cells.

Time cells across the multiCS sessions were classified into three distinct subtypes based on their reliability to specific trial types (figure 6C-F): sound-specific, light-specific, and modality-agnostic cells that respond to both conditions (169 sound-specific cells, 189 light-specific cells, and 88 modality-agnostic cells). Sound and light-specific time cells exhibited a preference for their respective CS modalities, encoding both the timing and identity of the CS, while modality-agnostic cells appeared to encode purely temporal information without CS specificity. When analyzing the peak times, sound and light-specific time cells did not retain their temporal firing patterns between sound and light trials (figure 6E-H). In contrast, modality-agnostic cells maintained consistent time signatures between both trial types, suggesting stable temporal encoding irrespective of modality (figure 6G). This pattern was evident regardless of whether the modality-agnostic cells were sorted by their peak times in sound trials (figure 6Ei,Fi)or light trials (figure 6Eii,Fii), emphasizing their stable time representation across conditions.

The classification of time cells into these subtypes was based on their CS selectivity, with a score of -1 indicating complete preference for light, 0 indicating equal firing across both modalities, and +1 indicating complete preference for sound. The average selectivity scores were 0.33±0.02 (Mean±SEM) for sound-specific time cells and -0.26±0.02 for light-specific time cells, with both groups showing significant deviation from 0 (WPT, p<0.0001). In contrast, the selectivity of modality-agnostic cells was not significantly different from 0, reinforcing their lack of CS preference. Figure 6I illustrates the multidimensional properties of individual time cells, plotting CS selectivity against peak firing time and time cell score (reflecting temporal precision). The three categories of time cells did not show a difference in temporal precision (p=0.59 Kruskal Wallis test) but interestingly, modality-agnostic time cells predominantly peaked during the stimulus or early post-stimulus period, while CS-specific time cells showed a more uniform distribution of peak times throughout the trial. This distribution was significantly different between modality-agnostic and modality-specific time cells (common vs. sound cells, p=0.00082; common vs. light cells, p=0.0000042; Kolmogrov Smirnov Test [KS]), but not between the sound- and light-specific groups (p=0.46), indicating that modality-agnostic time cells have a unique temporal firing pattern early in the trial.

In summary, some time cells in CA1 encode context, in the form of stimulus modality in TEC while others just keep time in a modality-agnostic manner. These subsets do not differ in their degree of time tuning but the modality-agnostic “pure” time cells have peak activity in and around the stimulus period and this timing remains similar between the two modalities. The modality-specific “contextual” time cells keep firing well into the post-stimulus period.

## Discussion

Our study mapped the emergence and dynamics of time cells in the hippocampal CA1 region across multiple stages of TEC, incorporating two sensory modalities which were first presented in individual sessions, then interleaved within a session. Neither the proportion nor precision of time locking, measured by two independent measures, changed with learning. Furthermore, time cell formation was not significantly influenced by the stimulus modality. Interestingly, network reliability exhibited a transient spike aligned with crossing learning criterion, suggesting its potential as an indicator of learning onset. While most time cells remapped between days, those cells that did persist retained time-selectivity within their original stimulus or post-stimulus epoch. Finally, we observed that a population of cells exhibited both modality and time encoding following the US of TEC, but modality-agnostic time cells were mostly confined to within 0.5 seconds of the US. We speculate that the extended modality-and-time encoding cells provide a substrate for learning beyond current task timing demands.

### Time cells emerge differently in different behavioural contexts

Time cells have been investigated in relatively few behavioural contexts, and their properties are quite context-dependent. Table 1 lists some of the key differences in task demands and time cell responses between the common behaviours used to study temporal processing.

**Table 1.** Comparison between the behavioural paradigms commonly used for studying temporal processing, focusing on the various cognitive demands and reported neuronal dynamics.

| Distinguishing parameters of the task | Spatial alteration/Tread mill run (Pastalkova et al., 2008) | DNMS (MacDonald et al., 2013, 2011; Taxidis et al., 2020) | Multiday Trace eyeblink conditioning (current study) | Trace fear conditioning with CS discrimination (Ahmed et al. 2020) |
| --- | --- | --- | --- | --- |
| Action involved (purposeful/goal oriented) | yes | yes | yes | no |
| Involves making a choice | yes | yes | no | no |
| Requires timed action | no | no | yes | no |
| Time cells observed | yes | yes | yes | no |
| Time cells correlate with learning | N/A | yes | no | N/A |
| Time cells/ensembles are distinct depending on context | yes | yes | partly | yes |
| Time cells predict individual trial errors | yes | no | no | N/A |
| Time cell persistence over days | N/A | ~5-15% (changes with learning) | ~10% | N/A |

We observe that time cells emerge before the animals show any signs of learning (Figure 3C,Di-iv). This is in line with previous studies (Modi et al., 2014; Taxidis et al., 2020) which report time cells on day 1. However, we find that time cells were present at a consistent proportion throughout learning (Fig 3Di-iv), whereas most studies that investigate temporal processing either report the presence of time cells, with some of their properties correlating with learning (Modi et al., 2014; Taxidis et al., 2020; Ma et al., 2024) or alternatively, the complete absence of time cells altogether (Ahmed et al., 2020). This might be because TEC sits in the middle ground of cognitive demand, where the animal has to take an action (similar to DNMS and spatial alternation but unlike fear conditioning) but does not perform choice based decision making (unlike DNMS and spatial alternation)

A more direct comparison is with a report which used single day learning of TEC (Modi et al., 2014). The study used harsher stimuli (strong airpuff and 90dB CS; as compared to our 65dB) both of which contribute to stress. It has been shown that stress increases CA1 excitability and plasticity (De Kloet et al., 1999; Shors, 2001; Shors et al., 1992; Shors & Servatius, 1997; Weiss et al., 2005), leading to rapid learning and learning related increase in time cell numbers.

Day-to-day remapping of time cells differs between DNMS and TEC. In DNMS there is partial stability of time-encoding between sessions (Taxidis et al., 2020). Mau et al., 2018 (water reward task with time delay) also reported time cells populations which were stable across days. In both DNMS and water reward protocols there is a consistent stimulus-driven rule which the animal has to retain from day to day. In contrast, in TEC we find that time cell identity is not maintained from day to day (figure 4B). We speculate that this is because of different requirements for remembering the modality, since in our TEC task there is no stimulus-choice decision, and the timing of the response is independent of modality.

### Network Reliability Increases Transiently During Learning Although time cell Proportions Do Not Change

Previous observations (Kim et al., 1995) show the hippocampus’s transient involvement during TEC learning, subsequently diminishing in significance for task recall. Our observations corroborate this with a network readout of time cell activity, which has a transient (1-day) peak before animals reach 60% behavioural criterion (figure 5). This phenomenon mirrors cellular-level observations from past research (Moyer Jr. et al., 1996), which noted an increase in CA1 pyramidal cell excitability post-learning acquisition, peaking 24 hours after and returning to baseline within seven days. Similarly, enhanced granule cell excitability has been documented in the dentate gyrus (Miller et al., 2022), upstream of CA1, correlating with learning progression. This too suggests a broader neural adaptability mechanism during learning phases.

### Time cells in TEC encode both task structure and modality

A unique feature of our behavioural protocol was the use of two distinct CS modalities (sound and light) in alternating blocks within the same session (MultiCS sessions). This design allowed us to detect two categories of time cells: those that encode the temporal structure of the task in a modality-agnostic manner as well as those that encode CS modality and therefore context. Prior studies in other paradigms have found that time cells encode both task structure as well as context. Ma et al., 2024 reported representation of valence and not stimulus identity in CA1 during a CS discrimination task. Even when the stimuli were swapped, the same cluster of CA1 cells flipped to encode the new CS+ (which was earlier CS-). In contrast, several studies report that time cells remap with any protocol modification, indicating their role in encoding context. Any change in the initial stimuli in a DNMS task triggers a reshuffling of CA1 time cells (MacDonald et al., 2013; Taxidis et al., 2020), suggesting these cells encode not only time but also its relation to specific memories. Similar observations were made in time cells in the MEC where changing the duration of stimulus presentation triggered distinct time cell sequences (Bigus et al., 2024).

### Time-encoding cells are active post-stimulus despite no subsequent timing requirements

Our modality-agnostic and modality-specific categories of time cells (Figure 6) differed in one key aspect: the former encoded time in and near the stimulus period (CS,trace,US) while the modality-specific time cell sequences extended atleast 1.7 seconds after the stimulus period (figure 6I). 60% of our time-tuned cells (∼13.5% of the total number of recorded neurons) were active post-stimulus (figure 3Dii,iv and Fi,ii). To our knowledge this is the first account of post-stimulus time-encoding in the hippocampus, though similar delayed activity has been reported in cortex (Curtis and Lee, 2010; Frank and Brown, 2003; Fuster, 2001; Goldman-Rakic, 1995). This observation has parallels with two other studies which report hippocampal cell sequences in a stimulus-free interval, with no subsequent behavioural requirement. The first is the observation that about 5% of hippocampal neurons participate in internally occurring cell sequences, in the absence of any external triggers (Villette et al., 2015). Here these sequences occurred when a mouse ran on a treadmill. The interpretation was that such sequences spontaneously emerge throughout motor activity. The second related observation is that of preplay, in which pre-existing reliable cell sequences recur as part of stimulus-driven place-cell activity (Dragoi and Tonegawa, 2011). The authors interpret these pre-existing sequences as a template onto which salient event sequences are associated. The key difference with our work is that neither of these two studies had a stimulus trigger. We propose that our observed post-stimulus time-encoding cells should also be considered to be time cells, in the sense that they encode time following an event, even though there is no terminating event or fixed window in which they occur.

An existing theoretical framework for our observation of post-stimulus sequences is that of echo-state or liquid-state networks, in which a stimulus triggers sustained reverberatory activity in a recurrent network (Buonomano and Maass, 2009; Laje and Buonomano, 2013; Maas and Markam, 2004; Maas et al., 2002).

We speculate that modality-agnostic time cells may be specialized for encoding the temporal structure of our TEC task, and hence their utility ends soon after the US. However, the cells that encode both modality and time (which are more common) may provide the network with a substrate for subsequent associations. Hence there is a potential computational rationale for their presence as a timing signal for many seconds after the end of the TEC period.

## Author contributions

Conceptualization by SB,USB; Methodology by SB,USB; Software by SB; Validation by SB, HN; Formal analysis by SB; Investigation by SB, HN; Resources by USB; Data curation by SB, HN; Writing–original draft by SB,USB; Writing–review and editing by SB,USB, HN; Visualization by SB, HN; Supervision by USB; Project administration by USB; Funding acquisition by USB.

## Acknowledgements

Funding support was from Department of Biotechnology grant BT/PR12255/MED/122/8/2016 and NCBS-TIFR core funding from Department of Atomic Energy, Government of India, Project Identification No. TRI 4006. We acknowledge NCBS campus facilities including Central Imaging and Flow Cytometry Facility, and Animal Care Resource Centre; and mechanical and electronic workshop. Anal Kumar assisted with GLM analysis to predict time from population and time cell activity and SVM analysis to classify CS modality based on neural activity. Dilawar Singh helped develop computer programs to run behaviour hardware.

## Methods

## Key Resources Table

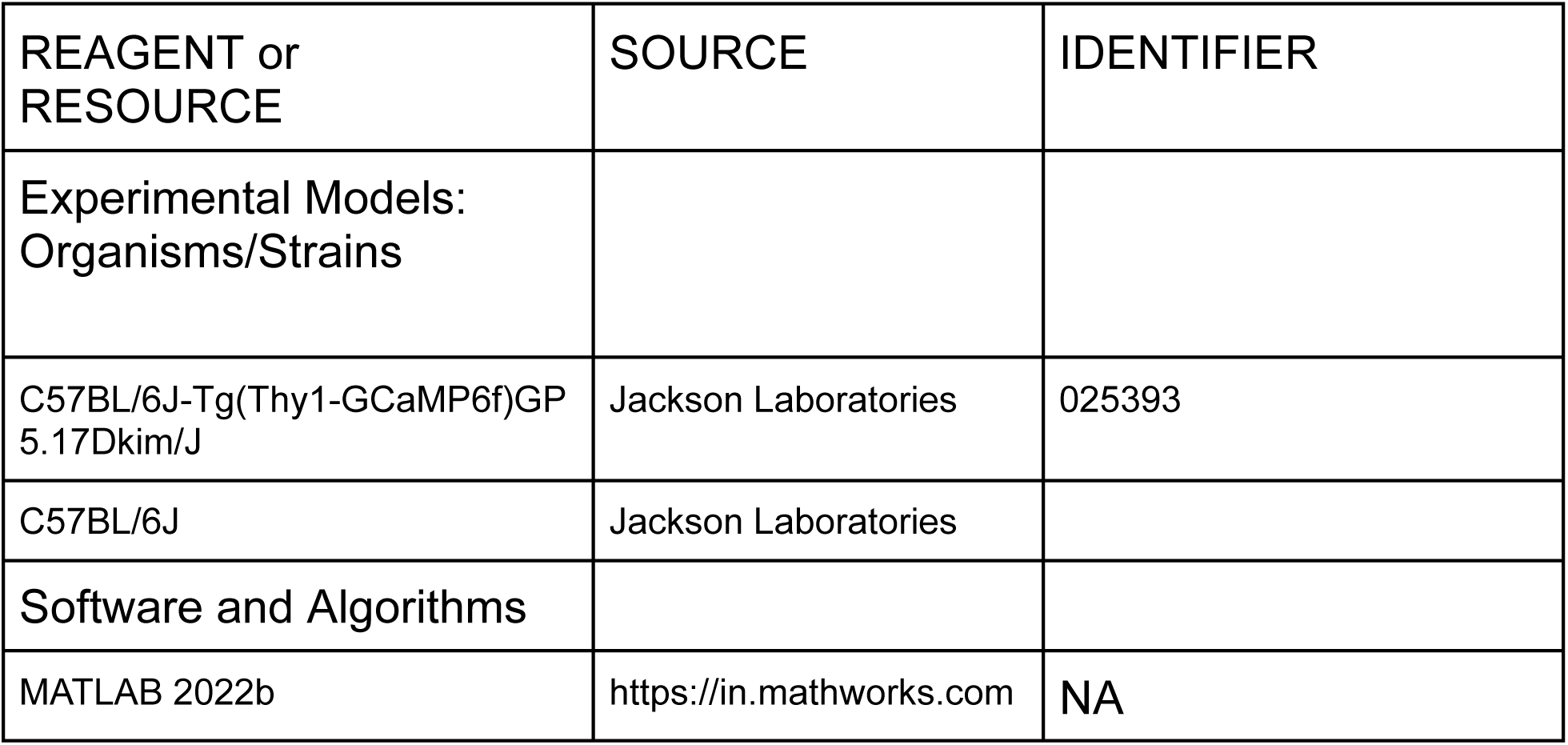
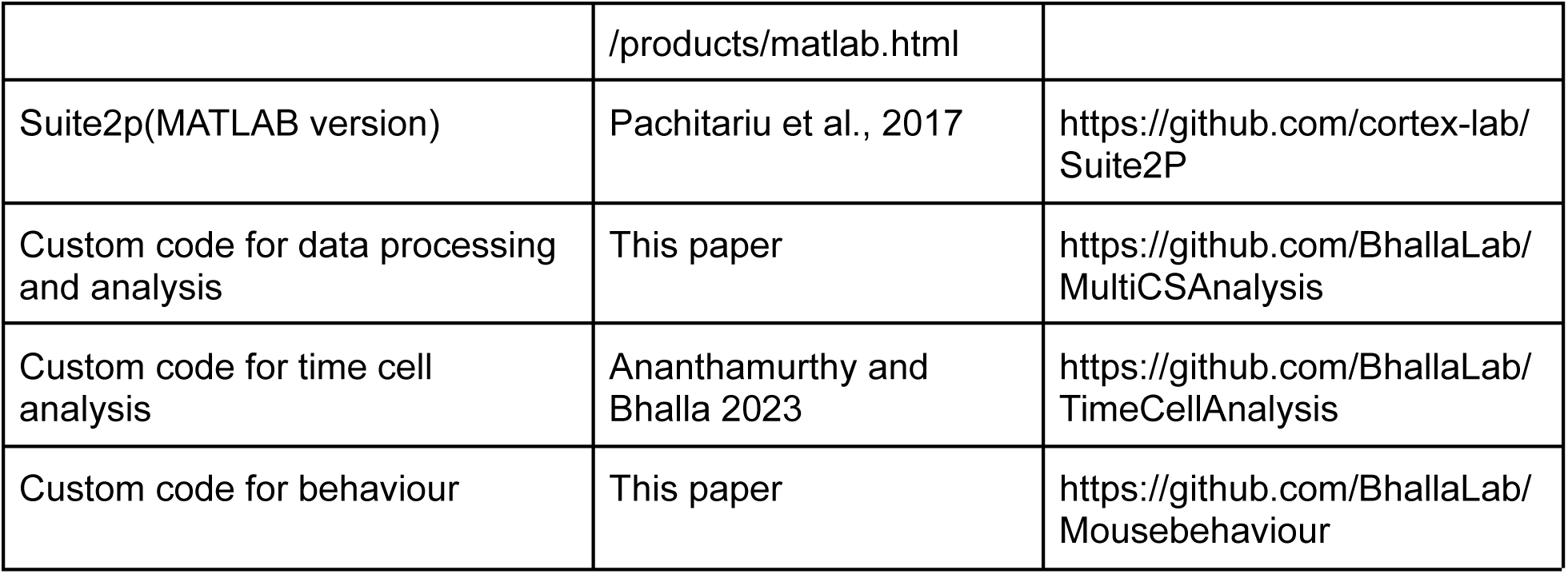

| REAGENT or RESOURCE | SOURCE | IDENTIFIER |
| --- | --- | --- |
| Experimental Models:<br>Organisms/Strains |  |  |
| C57BL/6J-Tg(Thy1-GCaMP6f)GP<br>5.17Dkim/J | Jackson Laboratories | 025393 |
| C57BL/6J | Jackson Laboratories |  |
| Software and Algorithms |  |  |
| MATLAB 2022b | https://in.mathworks.com | NA |
|  | /products/matlab.html |  |
| Suite2p(MATLAB version) | Pachitariu et al., 2017 | https://github.com/cortex-lab/Suite2P |
| Custom code for data processing and analysis | This paper | https://github.com/BhallaLab/MultiCSAnalysis |
| Custom code for time cell analysis | Ananthamurthy and Bhalla 2023 | https://github.com/BhallaLab/TimeCellAnalysis |
| Custom code for behaviour | This paper | https://github.com/BhallaLab/Mousebehaviour |

## List Of Abbreviations

TEC: Trace Eyeblink Conditioning
DNMS: Delayed Nonmatch to Sample
CA1: Hippocampus Cornu ammonis
CS: Conditioned Stimulus
US: Unconditioned Stimulus
pre: Pre-stimulus period
stim: Stimulus period (CS-trace-US)
Post and post-stim: Post-stimulus period
MW: Mann Whitney Test
F-Test: Fischer Test
OSWT: One Sample Wilcoxon Test
WPT: Wilcoxon Matched-Pairs Signed Rank Test
R2B: Ridge-to-Background
SEM: Standard Error of Mean
GLM: Generalized Linear Model
NP: Network Reliability Peak
HC: Day of hitting Behaviour Criteria (60%CR)
KWT: Kruskal Wallis Test
KS: Kolmogorov Smirnov Test
SVM: Support Vector Machine

## EXPERIMENTAL MODEL AND SUBJECT DETAILS

### Animals

A total of 10 GCaMP6f+/- (C57BL/6J-Tg(Thy1-GCaMP6f)GP5.17Dkim/J crossed with C57BL/6J), adult (12-24 week old) mice were used for in vivo two-photon calcium imaging experiments. An additional 9 mice which were GCaMP6f-/- and 12-24 week old were used for behaviour experiments without calcium imaging. All animals were experimentally naive. All animals were acquired from The Jackson’s Laboratory and were housed in cages of 2-4 animals, on a 12 hour light/dark cycle.

## METHODS DETAILS

### Surgical Procedures

To create the hippocampal window, we adopted the protocol previously reported by (Dombeck et al., 2010). All animals were water-deprived for 3-5 days before the day of the surgery till they reached 80% of initial body weight. Mice were anesthetized with Isoflurane (ISIFRANE 250, Abbott, North Chicago, IL, USA), vaporized and diluted with Carbogen (95% oxygen, 5% CO2) using a tabletop anesthesia machine and vaporizer (Item:901801 and 911103 from VetEquip Inc). 2 L/min vapor flow was used for induction and 1.2-1.5 L/min for keeping the mice under anesthesia while body temperature was maintained using a feedback-controlled heating pad (TC-1000 Temperature Controller and mouse heating pad from CWE Inc., USA). The mice were head-fixed using cheek clamps (Mouse Stereotaxic Adaptor, Stoelting Co.). Eye ointment (Chloramphenicol 1%w/w) was used to prevent desiccation of eyes when animals were under anesthesia. The fur on the animal’s head was trimmed using scissors until the scalp was cleanly exposed. The scalp was disinfected using 70% ethanol. A circular incision was made such that bregma, lamba, and about 5mm of the skull was exposed on either side of the sagittal suture. A cotton swab was used to remove fascia and to in general make the surface of the skull as dry as possible. Millimeters were marked on the skull using a marker to act as guides. A lightweight, custom-made stainless steel headbar with a circular central opening of 10mm diameter was attached to the skull using UV-curing dental cement (3M ESPE RelyX U200, Shade:TR). We did not use any skull screws for this process. Following this, a 3mm diameter burr hole was made using a dental drill. The burr hole was centered 1.5mm left lateral and 2mm rostral to bregma. The dura was removed using forceps, exposing the cortex underneath. A 26 gauge blunted needle (26×1/2 DISPOVAN syringes, HMD Ltd.) connected to a vacuum line was used to aspirate out the cortex, using constant washes with cortex buffer (125 mM NaCl, 5 mM KCl, 10 mM each of glucose and HEPES, 2 mM each of CaCl_2_ and MgCl_2_, and adjusted to pH 7.35 using NaOH) to prevent desiccation of the tissue. While the cortex buffer washes continually removed blood from the cavity, persistent bleeding if any was allowed to continue for 5-10 seconds as this helped the open blood vessel to clot. Cortical aspiration was done in a circular manner, leaving a cylindrical cavity of about 1 mm depth. This process took 15-20 mins. Aspiration was continued till the corpus callosum fibers were visible. The first two layers of the corpus callosum were removed leaving the third layer on the hippocampus. At this point, any remaining cortex buffer was suctioned out and the cavity was allowed to dry for 10-15 seconds till it lost the “glistening look”. A very small amount of Kwik-Sil (low-toxicity silicone adhesive from World Precision Instruments, Inc.) was applied directly to the exposed hippocampus. A stainless steel cannula which was previously prepared was slowly inserted into the cavity. The cannula had an outer diameter of 3mm and had a 3mm coverslip (D263 coverslip CS-3R, #0 thickness, Warner Instruments, LLC) attached at the lower end using a UV curing glue (Norland Optical Adhesive NOA 81). The coverslip sat directly on the hippocampus and the weight pushed the dab of Kwik-Sil to distribute it more evenly. The Kwik-Sil acted as a transparent adhesive between the cannula and the hippocampus and aided in reducing relative movement during imaging. Any gaps between the outer lateral surface of the cannula and the edge of the skull cavity were sealed with a second round of Kwik-Sil. Following this another round of dental cement was used to cover any exposed skull surface. Dental cement was made to flow around the lateral surface of the cannula and the upper part of the headbar for additional stability. This was done to reduce relative motion between the headbar, skull, and the cannula. The animals were then allowed to come out of anesthesia. Isoflurane was switched off and the animals were given a supply of Carbogen (5% CO_2_, 95% O_2_) at 1LPM till the breathing became normal. The animals were transferred to their home cage and kept at 1ml water per day till the end of the experimental period. They were given ibuprofen (2ml/L) and enrofloxacin (1ml/L) for 3 days ad libitum in water..

### Experimental Setup

A cylindrical foam cylinder(15.24 cm diameter, 11.43 cm length) with a metal axle through the axis which allowed 1D movement (forward and backward) was used as a treadmill. The previously implanted headbars were used to headfix the animals with the help of a custom-made clamp. Experiments were carried out in the dark in a chamber made of black, anodized steel which also housed the microscope. The microscope used was an in-house, custom-built, two-photon microscope. A Blue LED (480nm) positioned 7mm away and slightly towards the left of the mouse’s snout was used to deliver the light stimulus. Two speakers, one on each side and about 45cm away from the mouse were used to deliver a 3500Hz sound stimulus at 65dB. A wall-mounted supply of carbogen (∼15 psi), passed through a flowmeter (∼0.2 to 0.3 LPM, Cole Parmer) was used as an air source for the US. A PVC clear tubing with a 18 gauge syringe attached to the front was used to deliver the air-puff to the eye. The puff port was placed 5mm away from the animal’s left eye. The airflow was controlled using a solenoid valve (EV mouse valve, Clippard, Cincinnati, OH, USA). The eyeblink response was recorded at 200fps using a IR video camera (Blackfly S USB3; BFS-U3-13Y3M-C, Teledyne FLIR LLC) that was connected via USB to a laptop, where the data was saved.

The LED, speakers, solenoid, and camera were all controlled via an Arduino microcontroller which also provided a TTL pulse to trigger the imaging. The behaviour rig was controlled by a custom python software (https://github.com/BhallaLab/Mousebehaviour).

### Behaviour training and recording protocol

The Trace Eyeblink Conditioning protocol, adapted from Siegel et al., 2015, involved a 50 ms conditioned stimulus followed by a 250 ms trace interval and a 50 ms unconditioned stimulus, which was a puff of air directed at the animal’s eye. Our behaviour study comprised three stages of TEC termed Learning1, Learning2, and MultiCS.

Mice were handled for at least 3 days or until they were comfortable on the palm and did not jump off. The first day of handling coincided with the initiation of water restriction. Following the handling animals underwent craniotomy and headbar implant surgery. Mice were allowed to recover from the surgery for 5 days during which they were kept at a restricted water supply of 1ml/day. Their body weight was maintained at 80% or pre-water restriction weight.

The animals (n=9) were started on the CS1-US pairing, we called this learning1. Each day consisted of a single session of 60 trials(Figure 1Ai); out of which 10% were pseudo-randomly assigned as probe trials. The probe trials were CS only trials. 2 animals received sounds as CS1 and 7 received light. After the animals performed at 60% conditioned response criteria they were kept on the same protocol for 1.77±1.2 (mean±SD) days more for the response to stabilize (Supplementary Figure 1Bi). Animals (n=5) were then transferred to learning2 where the CS modality was switched; if it was light in learning1 then it was switched to sound, vice versa(Figure 1Aii). Learning2 continued till the animals hit the same criterion (Supplementary Figure 1Bii). Following this animals progressed to the multiCS stage where they received both pairings. Sound-puff and light-puff trials were presented in an interleaved manner, in blocks of 5 trials in each block (Figure 1Aiii). The last trial of every block was a probe trial. Animals (n=6) underwent multiCS sessions for 4.3±1.7 days (mean±SD). 1 out of the 6 animals considered for multiCS data also underwent both the previous protocols but weren’t included in learning1 and learning2 data due the animal not being naive before the first training.

### Behaviour-only Experiments

A cohort of 9 mice were headbar implanted but did not receive the craniotomy. These animals were used to give a better estimate of the behaviour learning curve but did not contribute to the calcium imaging data.

All 9 animals went through learning1. Out of them 4 animals received light as CS1 and were not moved to the next phases after they acquired learning1. Out of the remaining 5, 3 received light as CS1 and 2 animals got sound. The progression criteria remained the same as in calcium imaged animals-60% conditioned responses in a given session. These 5 progressed to learning2 where the first 3 underwent sound-puff pairings and the last 2 received light-puff pairings. All 5 animals were then moved to the multiCS phase where they got both sound-puff and light-puff trials, in blocks of 5 trials as mentioned in the previous section.

The mean learning curves of the behaviour-only animals were not statistically different from those that were coupled with imaging (KS test, p=0.7 for learning1 and p=0.97 for learning2). This allowed us to pool the dataset and create combined behaviour curves for learning1(n=18) and learning2(n=10) (Supplementary Figure 1Ci and ii).

### *In vivo* two-photon imaging

A custom-built two-photon microscope with galvo scanning (Model 6210H, Cambridge Technology, Inc.) was used to acquire calcium data, capturing approximately 100-150 cells within a 190 x 190 pixel field of view at a frame rate of about 11.5 Hz. The system utilized a Ti excitation laser (Chameleon Ultra II, Coherent) operated at 910 nm. GCaMP6f emission was collected through a water immersion objective (N16XLWD-PF - 16X Nikon CFI LWD Plan Fluorite Objective, 0.80 NA, 3.0 mm WD) and detected with an analog GaAsP PMT (H7422P-40; Hamamatsu, Japan). The amplified signal was binned for 2µs to construct each pixel. All imaging and behaviour experiments were conducted in complete darkness to prevent any light interference.

The imaging setup was controlled using LabVIEW 8.0 (National Instruments). To prevent the scan mirrors from producing auditory cues that could indicate the start and end of trials, they were kept active between trials for all but 2 animals. For G141 and G142 the scan mirrors were off during the inter trial interval. This may have cued the animals about the start of a new trial and we do not rule out a slight difference in neural response due to this. Imaging was synchronized with behavioural events using a TTL pulse from an Arduino microcontroller that also managed the behavioural setup.

## QUANTIFICATION AND STATISTICAL ANALYSIS

### Calcium Imaging Data Analysis

#### Preprocessing using Suite2p

Two-photon calcium imaging data were captured and saved in TIFF format using LabVIEW 8.0. These files were processed using the MATLAB version of Suite2p (Pachitariu et al., 2016), which performed automated ROI detection and motion correction.

Suite2p identifies ROIs by clustering pixels with highly correlated intensity profiles within a predefined size range of 10-15 um, approximately the size of a pyramidal neuron cell body. Each pixel within an ROI is weighted based on its correlation to the centroid intensity of that ROI, and the cell’s activity is calculated as the weighted sum of these pixels. Overlapping components and neuropil contamination was removed. ROIs were manually refined using Suite2p functions. Subsequent analyses were performed using custom scripts in MATLAB (R2022b).

#### dF/F calculation

The baseline fluorescence for each cell was calculated as the 10th percentile of the raw fluorescence trace for that trial. Fluorescence values for each frame were then baseline-subtracted, normalized, and converted into dF/F traces. These traces were stored as 3D matrices organized by cells, trials, and frames.

#### dF/F clean-up and filtering

In light-puff sessions, the microscope’s objective often captured an artifact from the LED flash used as the conditioned stimulus, which appeared in the neuronal data. To address this, we replaced the dF/F value at the CS frame with the median of the previous five frames. For consistency, we applied the same correction to sound-puff sessions as well.

Calcium transients from GCaMP6f typically span 180-200ms, corresponding to approximately 2-3 frames in our dataset. Signals shorter than this duration were identified as noise, often resulting from within-frame XY movement or Z-axis motion. To clean the data, we binarized the dF/F matrix, classifying values as signals only if they exceeded two standard deviations from the mean of that trial for each cell. This process effectively removed noise which typically manifested as single-frame signals.

#### Multiday Cell Registration

Cells were matched across days using a Suite2p function that facilitates the alignment of the same landmarks on different days. The fields of view are transformed and overlaid. An ROI is identified as the same cell across days if it overlaps by at least 60% with the ROI from the subsequent day. This process was extensively manually curated to eliminate any false positives.

Cells were classified as multiday time cells if they were detected on consecutive days and met the time cell criteria on each day. The percentage of multiday time cells was calculated based on the number of cells identified as time cells on a given day that could be matched to an ROI on the subsequent day.

## Time Cell Detection

Time cells, defined as neurons that fire consistently at the same time relative to the CS onset across trials, were identified using a custom C++ code (Ananthamurthy and Bhalla, 2023). This code employs two algorithms to detect time cells: the ridge to background method (Modi et al., 2014) and the temporal information method (Mau et al., 2018b).

For the ridge to background method, the peak response time (PT) of each cell was determined from the averaged dF/F traces spanning from the onset of the CS to 1.5 seconds after the US ended. This calculation used only alternate trials, while the remaining trials were employed to compute the ridge to background (R2B) ratio scores for assessing reliability. For these calculations, trials were averaged, and the summed area under the PT and its two adjacent points was determined. The ratio of this area to the area under all other points in the averaged trace was defined as the R2B ratio. For a control measure, these traces were randomly time-shifted before averaging. An independent PT was identified for each randomly shifted trace, and an R2B ratio was computed. This process was repeated 1,000 times for each cell’s data and the results were averaged. The R2B score for each cell was then calculated as the ratio of the R2B ratio for aligned traces to that for randomly shifted traces. Cells with R2B scores exceeding 2 were classified as time cells.

### Linear regression model to decode time from time cell activity

We employed a linear regression model to decode elapsed time from the CS onset using the neural activity of time cells. For each frame in a trial, starting from the first frame after the CS to the 24th frame after the CS, we constructed a neural activity vector which consisted of the dF/F of all the time cells in that frame. This resulted in 24 neural activity vectors per trial and n x 24 neural activity vectors per session where n is the number of trials in the session (typically 60). Each of these neural activity vectors were labeled by an integer corresponding to its frame number after the CS frame (0 to 23) excluding the CS frame. Some cells and frames were removed if any of their entries contained NaN and Inf. Scikit-learn’s LinearRegression was then trained to predict the frame number based on the neural activity vector. We report a 5-fold cross-validation score. The score is defined by:

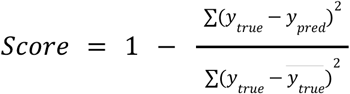

Where y_true_ it the true label of the neural activity vector, y_pred_ is the predicted label, and y̅_true_is the mean true label. If all the predictions match the true labels perfectly, the score is 1 while if the model performs worse than chance, the score can go below 0. Do note that while all y_true_ were positive integers ranging from 0 to 23, there were no explicit constraints imposed on the range or number type on y_pred_.

As control, we also shuffled the frame labels 1000 times for each session and trained a model on each such shuffle.

### Network Reliability

A network reliability parameter was defined to determine whether the network as a whole and not just individual cells was becoming more reliable with training. A reliable network would have a greater proportion of cells which are time tuned. Further, the degree of reliability of the individual cell would also have a positive correlation on network reliability. So, network reliability was defined as the sum of reliability scores of all time cells weighted by the proportion of time cells

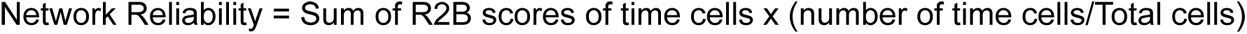

### Support vector machine for decoding CS identity

A Support Vector Machine (SVM) with linear kernel was employed to determine if cells encode information about the CS modality during multiCS sessions. A 2D neural-frame activity matrix was constructed for each trial which consisted of the dF/F of either all cells or only time cells across frames starting from the 1st frame after CS to the 25th frame after CS. This 2D matrix was rasterized to produce a single neural-frame activity vector. This vector was used to train an SVM with the labels corresponding to the CS used in that trial (0 for light-puff trials and 1 for sound-puff trials).Scikit-learn’s LinearSVC was used for this purpose. A 5-fold cross validation score was computed for each session which typically had 30 sound-puff trials and 30 light-puff trials. The score was defined as the ratio of correct predictions to total predictions. A perfect decoder would have the score of 1 while a decoder with no correct prediction would have a score of 0.

Additionally, as a control, the label for each neural-frame activity vector was shuffled 1000 times and scores were calculated. We also performed another negative control where singleCS light-puff sessions were used with each trial randomly labeled as light-puff or sound-puff. A decoder trained on this data would have by chance close to 50% accuracy.

### Behaviour Analysis

Eye blinks captured by the camera were saved as TIFF files. Using a custom MATLAB script, the eye region was identified and binarized into eye and non-eye areas. A central eye section was analyzed to determine eyeblink status, following the method described by Siegel et al. 2015. Each frame was assigned a Fraction Eye Closure (FEC) value, calculated by dividing the number of pixels representing the eye in that frame by the maximum number of pixels defining a fully opened eye in that session.

For each trial, a baseline was calculated by averaging the FEC values for 500ms before the conditioned stimulus (CS) onset. This baseline helped to minimize false positives, particularly if the animal’s eye was partially closed even before the CS was presented. An eyeblink response was defined as an FEC value exceeding 10% above the baseline. The formulas used were:

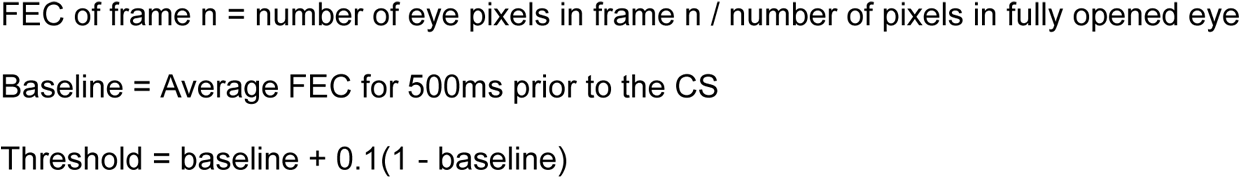

If the FEC value exceeded the threshold between the onset of the CS and the US, it was classified as a Conditioned Response (CR). To prevent false positives, trials in which animals exhibited excessive blinking during the pre-stimulus period were excluded if the Fano factor exceeded 0.5.

### Statistical analysis

All statistical tests were performed using GraphPad Prism 10. Test details are mentioned in the figure legends and main texts. Unless mentioned otherwise, nonparametric tests have been used as most distributions were not sufficiently close to normality as determined by Kolmogrov-Smirnov and Anderson-Darling tests (p>0.05).

## Supplementary Material

## Multimodal and interleaved trace-eyeblink conditioning

**Supplementary Figure 1:**
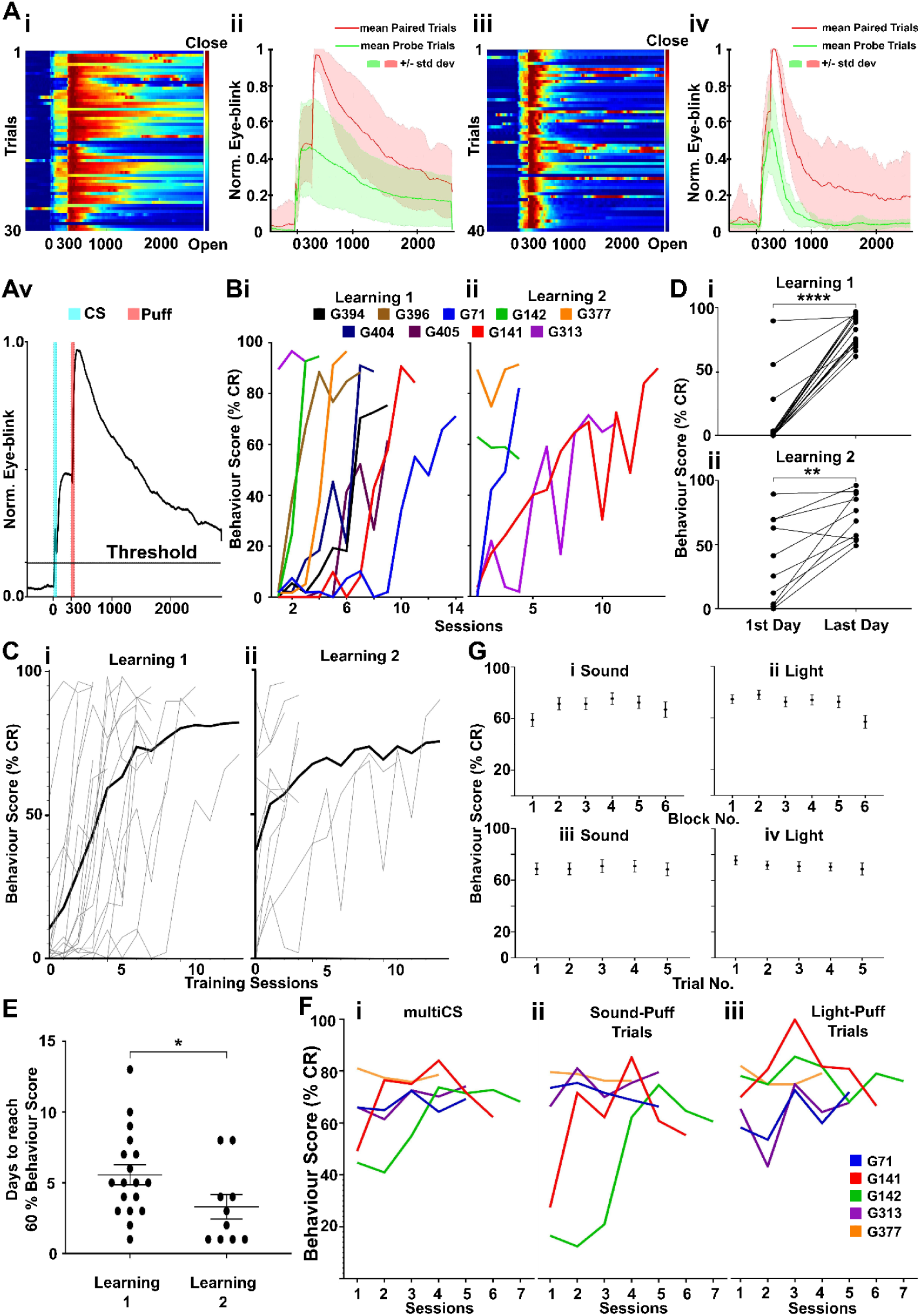
Three stage Trace Eyeblink Conditioning Responses and Learning dynamics. **A:** Post-learning eyeblink response for a multiCS session, showcasing conditioned response (CR) to both sound as well as light. In all cases the CS onset is at 500 ms and US onset at 800 ms. Ai: Heatmap for eye closure as a function of time for 60 sound-US trials. Aii: Normalized, trial average of Ai for paired (pink) and probe (green) trials. Aiii: Heatmap for eye closure as a function of time for 60 light-US trials. Aiv: Normalized, trial average of Aiii for paired (pink) and probe (green) trials. Av: Same as Aii but showing threshold for classifying eye closure as a conditioned response (Threshold = 10% of a full eye closure). **B**i: Learning curves for calcium imaged animals. Bi: Learning1: CS1-US, n=9 animals. Bii: Learning 2: CS2-US, n=5 animals. **Ci:** Learning curves for CS1-US of 18 animals (n=9 each for Calcium imaged and “behaviour only”). Animals were trained till they showed 60% CR and continued for an additional one to two days (1.77±1.2 days, mean±SD). **Cii** Learning curves for CS2-US of 10 animals (n=5 each for Calcium imaged and “behaviour only”). Gray lines show individual mouse’s data, black line shows the average. **D:** CR increases from the first to the last day of training, demonstrating successful conditioning acquisition. **Di**: data for learning1 (first day: 10.46±5.701, last day: 83.68±28.808. p<0.0001,WPT). **Dii**: Data for learning2 (first day: 37.61±10.56, last day: 72.2±5.687. p=0.005, WPT) demonstrating successful conditioning acquisition. **E:** Animals reach criterion quicker for learning2 compared to learning1. Time required to reach a 60% behaviour score for learning1 (5.55±0.715 days) is longer than for learning2 (3.3±0.869 days for 10 subjects), (p=0.04, MW). **F:** Animals perform at criterion for both stimulus modalities during interleaved trials. Fi: all trials. Fii: sound trials. Fiii: light trials. **G:** Consistent behaviour performance across all 5 trials in a block and all 6 blocks in a session, for each trial type in multiCS sessions. **Gi:** No significant differences are seen in animals’ performance across six blocks during sound-US (Kruskal-Wallis Test [KWT]).**Gii:** The light-US blocks showed slight difference,particularly in the block (p=0.02,KWT) but due to the differences being small we conclude that performance remains uniform. **Giii & iv:** behaviour performance remains uniform in all 5 trials within a block during multiCS sessions (sound-US: p-values=0.98; light-US: p=0.74. KWT), highlighting rapid adaptation to CS modality shifts.

All calcium imaged subjects (n=9) reached criteria (timed Conditioned Response on 60% or more of the trials in a session) for Learning1 on 5.1±1.1 days (mean±SEM)(Supplementary Figure 1Bi). Animals were still kept on the same protocol for 1 to 4 sessions more (1.7±1.2,mean±SD) before moving them onto the next phase, ie Learning2. Five of the initial nine animals reached this stage and they all reached criteria in 3.3±0.8 days(mean±SEM) (Supplementary Figure 1Bii). After this, in the multiCS stage, animals recalled the associations for both modalities from the 1st day onwards for 4.3±1.7 days (mean±SD).

To augment the dataset, additional “behaviour only” subjects, equipped with headbars but not subjected to craniotomy and cortical aspiration, were included in the analysis. We confirmed that the mean learning curves of the two sets were not statistically different (KS test, p=0.7 for learning1 and p=0.97 for learning2). 18 animals underwent learning1 (Supplementary Figure 1Ci), comprising both “imaging plus behaviour” (n=9) and “behaviour only” (n=9) groups. After the animals reached 60% Conditioned Response (CR) for learning1, they were kept on the same protocol for 1 to 4 additional sessions (1.77±1.2, mean±SD) to ensure that they had acquired the pairing before being moved onto learning2, and to obtain larger two-photon datasets for each learning condition. 10 animals underwent learning2 (Supplementary Figure 1Cii), evenly split between “imaging plus behaviour” and “behaviour only” groups.

There was an improvement in performance over the duration of both conditioning paradigms as measured by an increase in behaviour scores between the 1st and last day of training (Supplementary Figure 1Di and ii. Learning1 first day: 10.46±5.7, last day: 83.7±28.8, p<0.0001. Learning2 first day:37.61±10.56, last day:72.2±5.69, p<0.01. Mean±SEM, Wilcoxon matched-pairs signed rank test [WPT]).

The time to achieve a 60% CR rate decreased from Learning1 to Learning2, possibly indicating transfer learning effects from Learning1 (Harlow, 1949; Samborska et al., 2022; Tse et al., 2007). Learning1 took 5.55±0.7 days (n=18) while Learning2 took 3.3±0.87 days (n=10) to reach criterion (Supplementary Figure 1E. p<0.05 Mann Whitney test [MW])

The third stage was a multiCS phase where the animals had to recall both sound-US and light-US, in an interleaved manner in the same session. Animals performed near criteria for both modality even in early multiCS sessions (Supplementary Figure 1F. behaviour score for 1st multiCS session. Both modalities combined:75.62±6.559, sound-US:55.25±10.76, light-US:64.88±6.927; mean±SEM).

We found that performance across all six blocks within each modality remained consistent, Kruskal-Wallis Test (KWT) (p=0.24 and p=0.02) (Supplementary Figure 1Gi and ii). For light-US, performance across five trials in each block showed minimal differences.

We further tested whether animals required time to adjust while switching between blocks, since the first trial in each block was preceded by a trial with a different CS modality (Supplementary Figure 1Giii and iv) (sound-US p=0.98, light-US p=0.74 [KWT]). This finding demonstrates the animals’ ability to swiftly adapt to changes in CS modality without a significant performance drop.

### Network activity increases following stimuli in a learning-independent manner

**Supplementary Figure 2.**
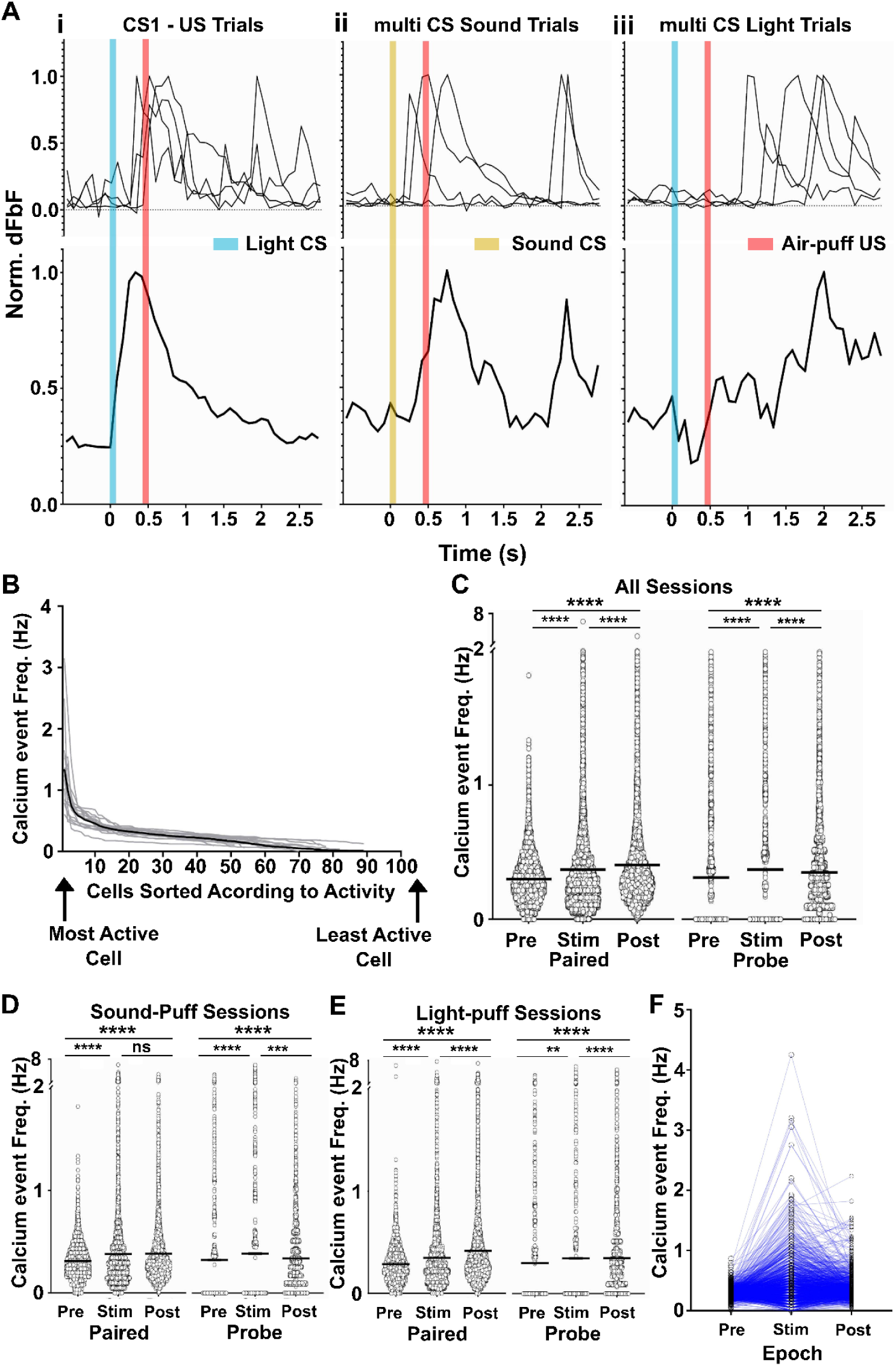
Stimulus-driven responses and learning effects on hippocampal CA1 network activity. **A:** Representative calcium activity from four trials of three exemplar cells across singleCS-US and multiCS sessions, illustrating individual trial (above) and trial-averaged (below) post-stimulus time histograms (PSTHs). **B**: Some cells show much higher activity than others. Cells arranged in descending order of activity (Calcium event frequency). Exemplar data from 15 sessions of mouse G142. Gray lines show individual sessions, solid black line shows the average. **C:** Aggregate calcium event frequen cy of frames in the 3 epochs. Data collected from across 133 singleCS-US sessions. Significant increases occur from pre-stimulus (pre=0.3±0.001 Hz) to stimulus (0.37±0.003 Hz) and post-stimulus periods (0.40±0.003 Hz). p<0.0001 (WPT) for all comparisons. Same observation held true for probe trials, indicating that the increase in activity wasn’t simply US driven. Pre=0.31±0.004 Hz, stimulus=0.37±0.006 Hz and post-stimulus=0.31±0.003 Hz. p<0.0001 (WPT) for all comparisons. **D, E**: Activity increases are not modality-specific. D: Sound-US: pre=0.3108±0.0028 Hz, stim=0.3798±0.0065 Hz, post=0.3826±0.0051 Hz; E: Light-US: pre=0.2867±0.003 Hz, stim=0.3474±0.0057 Hz, post=0.4156±0.0063 Hz; p<0.0001 for all comparisons, WPT). **F:** The active cell population is not constant in the 3 epochs. Individual cells change their activity states between epochs. Activity of cells in the pre-stimulus, stimulus and post-stimulus period as measured by calcium event frequency. Circles represent individual cells and lines connect the same cells in different epochs. Data Is from 1197 cells pooled from 19 sessions of G313

We next addressed the basic stimulus-driven responses of the pyramidal neurons in the hippocampal CA1 network activity. The hippocampus is known to receive multimodal sensory input (Acharya et al., 2016; Deshmukh & Bhalla, 2003; Ho et al., 2011; Komorowski et al., 2009; Liu & Otto, 2020). We divided each trial into 3 epochs: ∼1.7sec before the CS as pre-stimulus(pre), 350ms of CS, trace and US as stimulus(stim) and ∼1.7sec post US as post-stimulus(post). We defined calcium events as signals where the dfbf value was more than 2 standard deviations higher than the mean. Supplementary Figure 2A shows the calcium activity from 4 trials of 3 exemplar cells from a singleCS-US session and multiCS sessions(top row) and the trial averaged PSTH curves (bottom row). We tracked calcium activity in animals from naive to learned states and then in the recall phase. We looked at trial averaged activity from 133 sessions of 10,383 cells.

In every epoch we found that the total activity of the network was dominated by a small sub-population of cells, as would be expected since the hippocampus is known for sparse encoding (Jung & McNaughton, 1993; Skaggs et al., 1996) (Supplementary Figure 2B). Calcium event frequencies between the top 10% of active cells and the remaining cells across each observation period are shown in main text figure 2B. For each epoch (pre-stimulation, during stimulation, and post-stimulation) the top 10% active cells had markedly higher activity rates. This finding supports previous research, indicating that a small group of highly active cells, particularly in the Hippocampal CA1 region, are crucial in driving network responses. (Agarwal et al., 2014; Senzai & Buzsáki, 2017; Treves & Rolls, 1994).

We looked at trial averaged activity from 133 sessions over all stages of learning, from 10,383 cells and asked if stimulus delivery causes an increase in network activity. We found a statistically significant increase in network activity in the stimulus (0.37±0.003, p<0.0001. Mean ±SEM, WPT) and post-stimulus (0.4±0.003, p<0.0001. Mean ±SEM, WPT) epochs as compared to the pre period (0.3±0.001) (Supplementary Figure 2C). This increase was observed for probe trials as well which did not have a US. (pre=0.3±0.001, stim=0.37±0.003, post=0.4±0.003. p<0.0001 for both comparisons. Mean±SEM, WPT)

Does activity depend on whether the stimulus was a sound-US pairing or a light-US pairing? In both Sound-US (pre=0.3108±0.0028, stim=0.3798±0.0065, post=0.3826±0.0051; p<0.0001 for both pre vs stim and pre vs post WPT; Supplementary Figure 2D) and Light-US (pre=0.2867±0.003, stim=0.3474±0.0057, post=0.4156±0.0063; p<0.0001 for both pre vs stim and pre vs post WPT; Supplementary Figure 2E) pairing sessions, an increase in activity was observed during the stimulus and post-stimulus periods, indicating that the observed effect was not dependent on the stimulus modality. This led us to conclude that the network experiences a period of heightened activity following stimulus delivery, regardless of the stimulus type. Additionally, this increased activity was consistent in both paired and probe trials, indicating that the increase was not just due to the presence of an aversive US (p<0.0001 for all pre vs stim and pre vs post comparisons, p<0.001 for pre vs stim in light-US trials. WPT)

Supplementary Figure 2F illustrates that the same population of cells is not active in all 3 epochs. This is exemplar data from mouse G313, 19 sessions. The highly active cells during the stimulus phase were not active during the pre-stim period and many of them reduced their activity in the post-stimulus period.

### Time cell scores and proportions are independent of learning

**Supplementary Figure 3.**
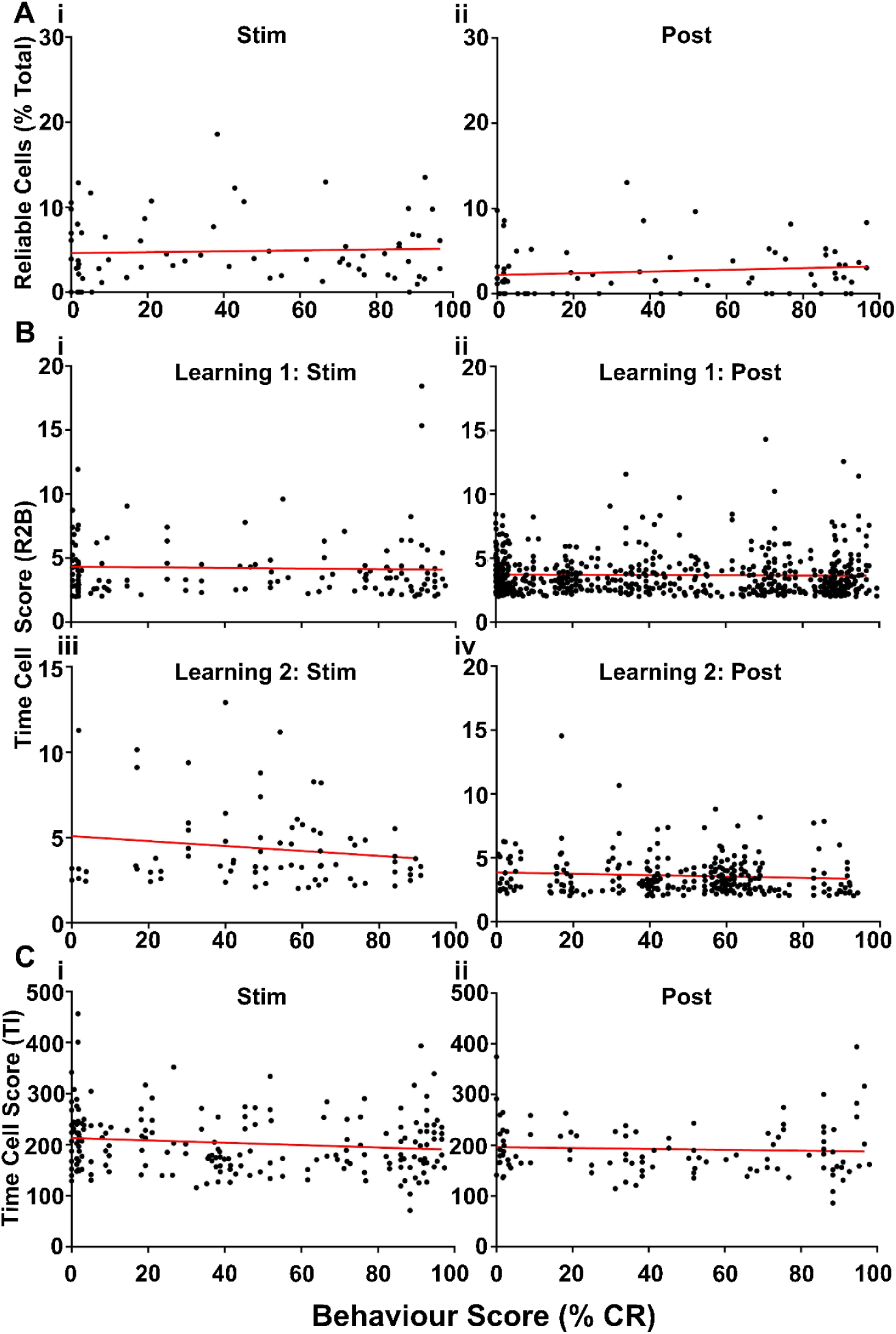
Time cell scores remain independent of learning progression according to both R2b and Temporal information method. Ai: Time cell percentage during stimulus = 4.8±0.45% for Learning1 (CS1-US) estimated using a temporal information (TI) metric (Mau et. al 2018). Aii: Post-stimulus percentage using TI metric=2.59±0.33%. The analysis confirms stable percentages across learning stages without significant correlation to behavioural scores (stimulus period slope: 0.005±0.012; post-stimulus period slope: 0.01±0.009; p=0.17 and 0.27 respectively, F-test). B: Cell-wise R2B score of time cells do not change over learning stages. Bi: R2B scores during the stimulus period of CS1-US show no significant changes across stages. Mean Score over all cells from all sessions during stimulus = 4.23±0.2. Linear regression analysis showed no correlation between R2B score and learning stage (regression line slope = -0.002±0.005; p=0.95, F-test). Bii: Mean Score during post-stimulus = 3.68±0.115 was agnostic of learning state as well (Regression line slope = -0.0008±0.003; p=0.8, F-test). Biii: Similarly for Learning2 (CS2-US), time cell scores during stimulus period = 4.35±0.272 did not correlate with behaviour performance (regression line slope = -0.014±0.001; p=0.41, F-test). Biv: Mean R2B score for the post-stimulus period of CS2-US was 3.59±0.09 and it remained unchanged during learning (slope= -0.005±0.003; p=0.82, F-test). C: Cross-validation with the temporal information method yields scores of Ci: 203.6±4.126 for the stimulus and Cii: 192.5±4.923 for the post-stimulus period, with no significant correlation to behavioural scores (slopes: -0.231±0.114 for stim and -0.08±0.142 for post; p=0.04,0.54, F-test), reinforcing the stability and independence of time cell distribution from learning progression.

Time cell proportions, as determined by R2B scores greater than 2, were independent of learning state. We verified this observation using three other methods. First was the temporal information metric (TI) outlined by Mau et al 2018. The second was the R2B score itself, as opposed to the proportion of cells above a certain R2B score.

Using TI, we identified 4.8±0.451% and 2.59±0.331% (mean±SEM) of cells as time cells for the stimulus and post-stimulus periods, respectively (Supplementary Figures 3 Ai and ii). Consistent with the initial method, the correlation between the proportion of time cells and the animals’ behavioural scores remained nonsignificant.

Next we checked if an analog readout of the extent of time-locking might be a more sensitive readout instead of a binary classification of time cells. To do this we looked at their individual R2B scores (Supplementary Figure 3 Bi-iv). This score reflects the extent of time-locking for each cell. The more precise a cell’s firing with respect to a timepoint and the more trials a cell fires in that time field, the higher is this score. Using this metric the dorsal-most CA1 were found to be agnostic of the animal’s behaviour performance, corroborating the result from the time cell proportions.

As a final validation we checked the TI scores (and not just the binarized %) of the cells Mau et al 2018. This metric showed the same result. The TI scores of cells did not correlate with the increase of behaviour scores, indicating that time cell behaviour remained unchanged throughout the learning process (Supplementary Figure 3 Ci and ii).

These three parallel methods corroborate the initial observations, affirming the stable presence and distribution of time cells across the learning stages, as determined by both direct measurement and temporal information analysis (Main figure 3 Di-iv).

### Time cell scores and proportions are independent of learning

**Supplementary Figure 4:**
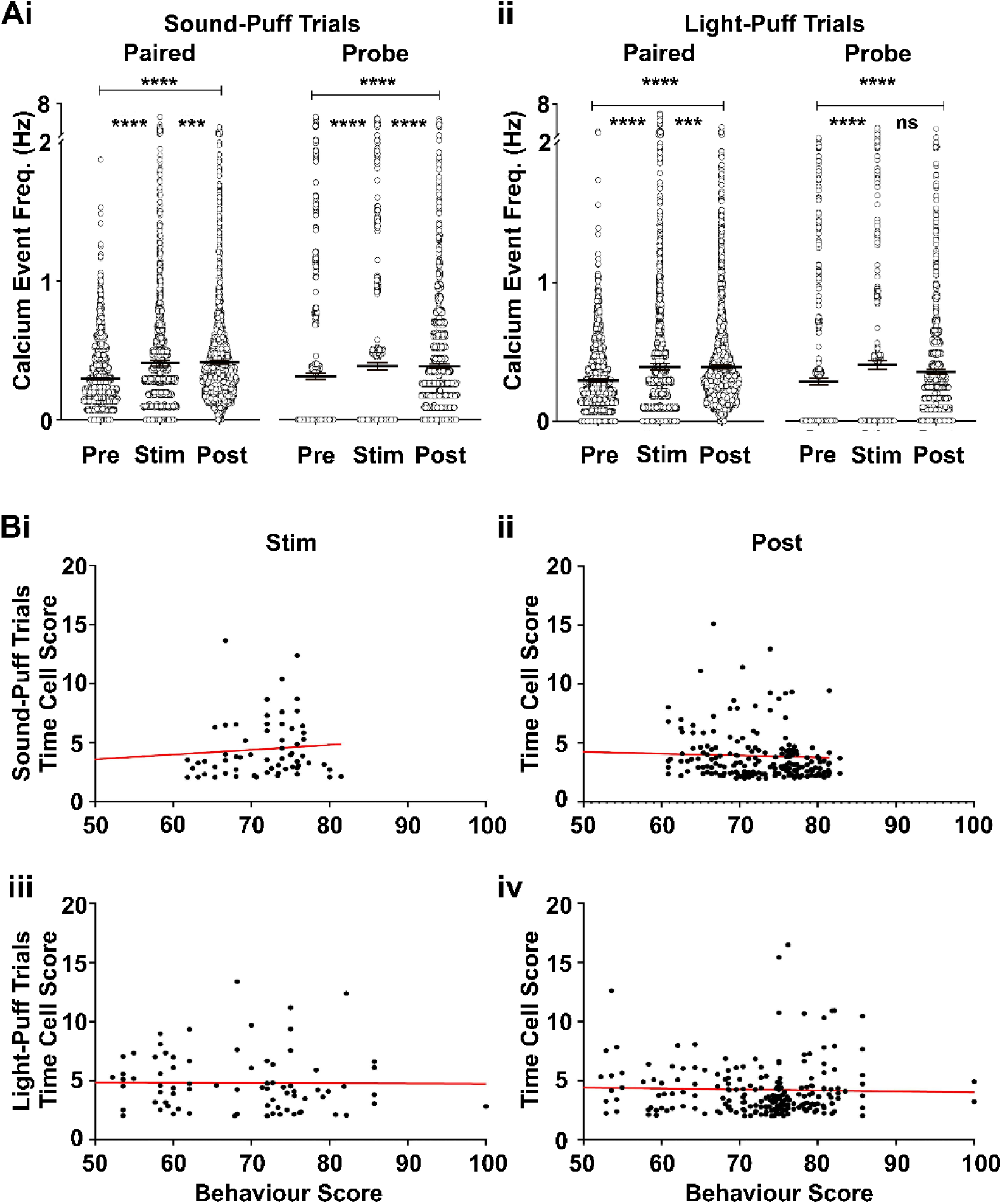
Neuronal Activity and Time Cell Dynamics Across Stimulus Modalities. Ai-ii: Increase in calcium event frequency in both sound-US and light-US trials due to stimulus delivery. Each point is trial averaged activity in a frame which falls under one of the epochs. For sound-US when going from pre-stimulus (0.29±0.005 Hz) to stimulus (0.4±0.009 Hz) and post-stimulus periods (0.41±0.007 Hz) there is significant increase in neuronal activity (p<0.0001, Wilcoxon test). This also holds true for light-US; pre-stimulus (0.29±0.005 Hz) to stimulus (0.39±0.01 Hz) and post-stimulus periods (0.39±0.007 Hz) (p<0.0001, Wilcoxon test). Same trends are visible in paired as well as probe trials. Bi-iv: Linear regression analyses of time cell activity versus behavioural scores for sound-US (i, ii) and light-US modalities (iii, iv), for stimulus periods (left) and post-stimulus (right). Sound-US trials show slopes of 0.04±0.029 (stim) and -0.015±0.013 (post), while light-US trials present slopes of -0.002±0.022 (stim) and -0.008±0.12 (post), with p-values (0.17, 0.24, 0.92, 0.52 respectively for the four groups) indicating no significant correlation across modalities (F-test).

We found stimulus-dependent increase in activity in both sound-US and light-US trials(p<0.0001 [WPT]; figure 6Ai and ii) in both the stimulus and post-stimulus periods compared to the pre-stimulus period. Additionally, this increased activity was consistent in both paired and probe trials, indicating that the increase was not just due to the presence of an aversive US (p<0.0001 for all pre vs stim and pre vs post comparisons, p<0.001 for pre vs stim in light-US trials. WPT). Thus the occurrence of both CS and US consistently elevated network activity, irrespective of the trial modality.

We then asked whether the activity of time cells correlated with behavioural scores. We found that time cell scores were independent of the behavioural scores for both modalities (figure 6Bi-iv). The slope of the linear regression line for sound-US trials during the stimulus phase was 0.04±0.029, and for the post-stimulus phase, -0.015±0.013. For light-US trials, the stimulus phase slope was -0.002±0.022, and for the post-stimulus phase, -0.008±0.12. These values are not significantly different from zero (F-test), indicating no correlation between time cell score and behaviour score.

